# High-resolution low-cost LCD 3D printing of microfluidics

**DOI:** 10.1101/2023.12.31.573772

**Authors:** Houda Shafique, Vahid Karamzadeh, Geunyong Kim, Yonatan Morocz, Ahmad Sohrabi-Kashani, Molly L. Shen, David Juncker

## Abstract

The fabrication of microfluidic devices has progressed from cleanroom manufacturing to replica molding in polymers, and more recently to direct manufacturing by subtractive (e.g., laser machining) and additive (e.g., 3D printing) techniques, notably digital light processing (DLP) photopolymerization. However, many methods require technical expertise and while DLP 3D printers remain expensive at a cost ∼15-30K USD with ∼8M pixels that are 25-40 µm in size. Here, we introduce (i) the use of low-cost (∼150-600 USD) liquid crystal display (LCD) photopolymerization 3D printing with ∼8M-58M pixels that are 18-35 µm in size for direct microfluidic device fabrication and (ii) a poly(ethylene glycol) diacrylate-based ink developed for LCD 3D printing (PLInk). We optimized PLInk for high resolution, fast 3D printing and biocompatibility while considering the illumination inhomogeneity and low power density of LCD 3D printers. We made lateral features as small as 75 µm, 22-µm-thick embedded membranes, and circular channels with a 110 µm radius. We 3D printed microfluidic devices previously manufactured by other methods, including an embedded 3D micromixer, a membrane microvalve, and an autonomous capillaric circuit (CC) deployed for interferon-γ detection with excellent performance (limit of detection: 12 pg mL^-1^, CV: 6.8%), and we demonstrated compatibility with cell culture. Finally, large area manufacturing was illustrated by printing 42 CCs with embedded microchannels in <45 min. LCD 3D printing together with tailored inks pave the way for democratizing access to high-resolution manufacturing of ready-to-use microfluidic devices by anyone, anywhere.

## Introduction

Microfluidics, through miniaturization in micrometer-sized vessels and microchannels, can reduce the fluid volumes required for analysis and synthesis to microliters and less, form the foundation for lab-on-a-chip devices, and are amenable to automation.^1, 2^ However, wider adoption of microfluidics and lab-on-a-chip devices in diagnostics, synthesis, and research is slowed by complex fabrication processes. Microfluidics emerged as a field thanks to cleanroom microfabrication inherited from the semiconductor industry relying on photolithography and using silicon or glass microfabrication methods which are dependent on capital cost intensive equipment. Soft lithography methods helped relieve the dependency on the cleanroom as multiple replicates from a single microfabricated mold could be made in a common research lab, and greatly accelerating the adoption and dissemination of microfluidics for primarily research applications.^3^ More recently, direct manufacturing methods have been introduced including subtractive ones such as laser ablation^4^ or micromilling,^5^ but offer limited relief due to drawbacks such as the need for expensive equipment, technical expertise, or provide limited resolution.

Additive manufacturing, and in particular 3D stereolithography (SLA) printing characterized by layer-by-layer UV patterning and photopolymerization of successive layers in a photocurable ink to build up a 3D printed object, has received considerable attention thanks to its affordability, high-resolution, and ease-of-use.^6, 7^ A layer is exposed to a digital pattern that solidifies the ink within a defined layer thickness; the layer then rises to allow uncured ink to fill the void, followed by digital photopolymerization of the new layer, and the process repeats iteratively. In an effort to clarify the terminology, we distinguish three methods of SLA and strategies to selectively expose ink within the layer: (i) laser scanning SLA operating with a galvanometer, (ii) digital light processing (DLP-SLA) that relies on a digital micromirror device and an optical system for projecting a pattern, and most recently (iii) masked SLA using a liquid crystal display (LCD) 3D printer where collimated light is directed through an LCD screen that digitally renders the design and photopolymerizes ink atop the LCD.

Laser SLA gained popularity thanks to high-resolution prototyping on a large print bed (335 × 200 × 300 mm^3^) for microchannels ranging between 250-500 µm with 30-140 µm laser spot sizes.^8^ Low-force SLA using a flexible vat reduces the adhesion force between formed layers and the bottom of the vat for intricate microfeature formation (e.g., separation membranes).^9^ Additionally, many materials used in laser SLA are biocompatible,^10^ but have largely been limited to commercial inks with proprietary formulations. Further, the single spot photopolymerization process with one or two lasers increases build times, especially for microfluidic devices that are generally blocks of solid ink with few voids that constitute the channels.

DLP 3D printing became widely adopted for microfluidics thanks to rapid and high-resolution fabrication with reported microchannels as small as 18 × 20 µm^2^, 3D printer pixel sizes ranging from 2-40 µm, and an illumination wavelength between 365-405 nm that can be used to photocure a wide range of materials.^11–13^ The availability of open-source printers, online design repositories (e.g., Thingiverse, GrabCAD, Printables), tailored workflows (e.g., print-pause-print for multimaterial designs),^14^ and custom ink formulations further increase the potential. The development of open-source inks such as those based on poly(ethylene glycol) diacrylate (PEGDA) for DLP 3D printing benefit from known compositions, which could help evaluate the impact of leachable and washable cytotoxic photosensitive components, and can be tailored and optimized for high-resolution embedded 3D printing, enhanced mechanical properties, low viscosity for fast printing, as well as for low protein adsorption and cytocompatibility.^7, 15, 16^

Altogether, the synergy of high-precision 3D printers, custom inks, and direct 3D printable designs enables digital manufacturing, i.e., the seamless and automated fabrication from digital file to final product with minimal post-processing. Digital manufacturing of microfluidic components has been possible early on, and now extended to the fabrication of fully functional systems based on capillary flow.^7^ Indeed, as capillary microfluidics can operate without peripherals,^17^ and complex fluidic algorithms could be structurally encoded into so-called capillaric circuits (CCs),^18, 19^ our group showed digital manufacturing functional systems in the form of CCs. Thanks to custom intrinsically hydrophilic inks, ready-to-use CCs systems, could thus be printed using DLP 3D printers.

However, the capital cost of common research-grade microfluidic DLP 3D printers (∼15K-30K USD) constitute a significant entry barrier for many potential users. Furthermore, while the pixel numbers have increased, with many printers culminating at 3840 × 2160 ≅ 8M pixels, the trade-off between print resolution and build area has not been resolved for microfluidics which requires small pixel size, and hence small build areas, but come at the cost of limited manufacturing throughput.

LCD photopolymerization 3D printers retail for as little as ∼150-600 USD, with pixel numbers of 4K (>8M pixels), 8K (>33M pixels), and up to 12K (>58M pixels), and pixel size of 18-50 µm, thus outperforming DLP 3D printers both in terms of number of pixels and affordability. LCD 3D printers utilize an array of discrete light-emitting diodes (LEDs) that can now be mounted at high density (i.e., chip-on-board, COB) and that are collimated by an optical system (e.g., COB lens and Fresnel lens) then pass through an LCD screen to reach the vat bottom. The number of pixels has been growing exponentially, and with a range of pixel sizes that extend to smaller dimensions, thus offering both higher density and larger print areas, and the capacity to print high resolution structures such microfluidics on large print beds. However, in a recent study, Caplins *et al.* report illumination non-uniformity due to variable irradiance and spectral differences in discrete LEDs resulting in inconsistent prints.^20^ Furthermore, the 50% transmittance loss of LCD screens by the crossed polarizers further reduces the irradiance of LCD 3D printers (2-3 mW cm^-2^) compared to their DLP counterparts (5-100 mW cm^-2^). Printing more voxels per time requires higher irradiance as the rate of printing for a given ink is limited by the power density of the light source.^21^ Lastly, LCD screens degrade rapidly at low wavelengths and are thus limited to >400 nm illumination, which reduces material selection and ink efficiency.^12, 20, 22, 23^ Prior work has shown success in leveraging LCD 3D printing for microfluidic master mold fabrication,^24–27^ but the potential for throughput manufacturing on large build plates and direct LCD 3D printing of open and embedded microchannels has not been shown.

Here, we present high-resolution fabrication of embedded and open microfluidic devices using low-cost LCD 3D printing with a custom formulated low-viscosity PEGDA-based ink that cures using low irradiance and minimizes the effect of illumination variability on curing depth. The lateral and vertical resolution of open and embedded structures are characterized using a series of test structures, and showcases high fidelity and dimensionally accurate printing of open and embedded structures down to a resolution in the tens of micrometers. The biocompatibility of the ink is validated based on an ISO standard for cell toxicity. Three microfluidic devices are manufactured by LCD 3D printing and characterized: (1) a microfluidic mixer previously made by laser micromachining, (2) membrane microvalves commonly made by replica molding, and (3) CCs previously made by DLP 3D printing. To illustrate the advantages of LCD 3D printers with high pixel numbers, we manufacture 42 CCs at once in <45 min showcasing both large area printing and potential for high throughput manufacturing.

## Materials and Methods

### Materials

#### Ink materials

Poly(ethylene glycol) diacrylate (PEGDA)-250, Cat. #475629, lot #MKCS0146, Sigma-Aldrich, Oakville, Ontario, Canada); diphenyl(2,4,6-trimethylbenzoyl)phosphine oxide (TPO), Cat. #415952, lot #MKCK2346, Sigma-Aldrich, Oakville, Ontario, Canada), 2-isopropylthioxanthone (ITX) (Cat. #I067825G, lot #ZNNQE-KT, TCI America, Portland, Oregon, United States); pentaerythritol tetraacrylate (PETTA) (Cat. #408263, lot #MKCR5556, Sigma-Aldrich, Oakville, Ontario, Canada).

#### Other chemicals

3-(Trimethoxysilyl)propyl methacrylate (Cat. #M6514, lot #SHBG7600V, Sigma-Aldrich, Oakville, Ontario, Canada; fluorescein sodium salt (Cat. #46960, lot #2082530, Sigma-Aldrich, Oakville, Ontario, Canada), isopropyl alcohol (IPA) (Fisher Scientific, Saint-Laurent, Quebec, Canada).

#### Immunoassay

Purified mouse monoclonal IgG anti-human interferon-γ capture antibody (Cat. #MAB2852, lot #FIO1022021, R&D Systems, Minneapolis, Minnesota, United States), biotinylated affinity purified goat IgG anti-human interferon-γ detection antibody (Cat. #BAF285, lot #ZX2721071, R&D Systems, Minneapolis, Minnesota, United States), recombinant human interferon-γ protein (Cat. #285-IF, lot #RAX2422031, R&D Systems, Minneapolis, Minnesota, United States), Pierce streptavidin poly-horseradish peroxidase (pHRP) (Cat. #21140, lot #XJ360080, Thermo Fisher Scientific, Waltham, Massachusetts, United States), SIGMAFAST 3,3’-Diaminobenzidine tablets (Cat. #D4293, lot #SLCG5357, Sigma-Aldrich, Oakville, Ontario, Canada), bovine serum albumin (BSA) (Cat. #001-000-162, lot #162191, Jackson ImmunoResearch Labs, West Grove, Pennsylvania, United States), BSA-biotin (Cat. #A8549, Sigma-Aldrich, Oakville, Ontario, Canada), Tween 20 (Cat. #P7949, lot #SLBX0835, Sigma-Aldrich, Oakville, Ontario, Canada).

All assay reagents were prepared using 1X phosphate-buffered saline (PBS) (pH ∼ 7.4) supplemented with 0.05% Tween 20 and 5% BSA. All other solutions were prepared using water from a Milli-Q system (resistivity: 18 MΩ cm; Millipore).

### Ink preparation

The 3D printing ink was based on a low molecular weight PEGDA-250 supplemented with 0.5% (wt/wt) diphenyl(2,4,6-trimethylbenzoyl)phosphine oxide (TPO) photoinitiator, 1.5% (wt/wt) 2-isopropylthioxanthone (ITX) photoabsorber, and 2% (wt/wt) pentaerythritol tetraacrylate (PETTA) crosslinker. The reagents were mixed in a 500-mL amber glass bottle under magnetic stir for at least 2 h before use and stored at room temperature thereafter.

### 3D printing of microfluidic chips

The microfluidic chips were designed either in AutoCAD (Autodesk) or Fusion 360 (Autodesk), then exported as an STL file for slicing in a third-party software, CHITUBOX, at a layer thickness of 20 µm. The slices were uploaded to the Elegoo Mars 3 Pro, Elegoo Mars 4 Ultra, or Elegoo Saturn 2 (ELEGOO, Shenzhen, China) masked stereolithography LCD 3D printers with a 405 nm light source. Print settings for all the devices presented here are given in **Table S1**. The printed chips were washed on the build plate to remove excess uncured resin with IPA and dried with compressed air or nitrogen, followed by 1 min of UV curing (CureZone, Creative CADWorks, Concord, Ontario, Canada). Embedded devices were ready for use following UV curing; meanwhile, open channel CCs were sealed with a pressure adhesive tape (9795R microfluidic tape, 3M, Perth, Ontario, Canada) to encapsulate the microchannels.

### Cure depth, light penetration depth, and absorbance measurements

To determine the penetration depth of light, 50 × 75 × 1 mm^3^ glass slides were first cleaned with IPA, then silanized via liquid phase deposition by immersing a glass slides in a solution of 2% 3-(trimethoxysilyl)propyl methacrylate prepared in toluene for at least 2 h or overnight. The slides were then cleaned in fresh toluene and dried with compressed nitrogen. The treated glass slides were placed directly on the 3D printer LCD screen; with the UV illumination on, the power intensity was read through the glass using a UV light meter with a 405 nm probe (Model 222, G&R Labs, Santa Carla, California, United States) to be 2.23 mW cm^-2^. Then, 8 µL of uncured ink was placed on the glass slide, and the UV light was illuminated at different exposure times and repeated for each ink formulation. Following exposure, the glass was cleaned with IPA to remove excess uncured ink, dried with compressed nitrogen, and the cure depth of the formulation was measured using a stylus profilometer (DektakXT, Bruker, Billerica, Massachusetts, United States) that was configured to measure using a 12.5 µm probe radius with a 3 mg force to scan a 7 mm region in 20 s. The cure depths were recorded in Vision64 and the average height was measured according to the International Organization for Standardization (ISO) 4287 protocol after 2-point leveling to record the baseline.

The light absorbance of each photocurable ink was measured using a spectrophotometer (NanoDrop@ND-1000, NanoDrop Technologies, Wilmington, Delaware, United States). A blank reading was performed using MilliQ water, followed by recording the light absorption spectra with 2 µL of ink solution at a 0.1 mm path length.

### Cell culture and cytotoxicity assay

To assess the cytotoxicity of the ink formulation, a cytocompatibility assay was performed in compliance with ISO 10993-5:2009 standards. The cells used in this study were kindly provided by Dr. Arnold Hayer of McGill University,^28^ and they were grown and passaged according to ATCC’s recommendations and cultured in EGM-2 media. Briefly, 8 × 3 mm^2^ (diameter × thickness) rings were 3D printed and washed for 72 h with 70% ethanol with daily refresh of ethanol and then washed with PBS for 48 h to remove any unreacted photoactive components. The rings were then co-cultured with mCherry-labelled human umbilical vein endothelial cells (HUVECs) seeded at a density of 10,000 cells per well. Quantitative cell viability measurements were performed every 24 h over a total of 72 h using the PrestoBlue^TM^ cell viability reagent. HUVECs seeded at an identical density were cultured alongside the ring co-culture as a control and used to establish 100% cell viability for each time point. Both the control and co-culture conditions were imaged every 24 h over a total of 72 h using a Ti2 inverted microscope and analyzed using NIS-Element (Nikon, Japan) for all biological replicates.

### Numerical simulation of concentration fields in the micromixer

The concentration field of the micromixer was calculated by the finite element method using COMSOL Multiphysics v.5.6 (COMSOL, Inc., Burlington, Massachusetts, United States). The diffusion coefficient of fluorescein (4.25 × 10^-10^ m^2^ s^-1^) was applied to solve the steady-state concentration field of fluorescein at a flow rate of 0.1 mL min^-1^. The concentration field was sliced into cross-sections to obtain the splitting and recombining stream profiles along the length of a mixing unit.

### Fluidic demonstrations

To visually assess the fluid flow in the microfluidic chips, a 2% solution of food dye in MilliQ water was prepared and loaded in the chips. For the micromixer, a 10 µM solution of fluorescein was prepared in MilliQ water. In the case of the ELISA-chips, the devices were assessed with a solution of 2% food dye in 1X PBS buffer containing 0.05% Tween 20.

### Fluorescent imaging through microfluidic chips

To facilitate microscopy imaging of fluorescent solutions in the chips, micromixer devices were mounted to a glass slide by UV photopolymerizing a drop of uncured resin between the chips and a plain glass slide (25 × 75 × 1 mm^3^) for 40 s. The device was printed with cylindrical ports connected to a programmable syringe pump (Kd Scientific KDS250) via Tygon E-3603 tubing to flow solutions into the micromixer at known flow rates. Fluorescent images were acquired using a Nikon Ti2 inverted fluorescence microscope using NIS elements. Flow profiles were analyzed in ImageJ2 Ver. 2.9.0/1.53t (public domain software, National Institute of Health, Bethesda, Maryland, United States).

### Flow rate measurements through the microvalve

To assess the functionality of the microvalve, black dyed water was flown continuously through the flow channel inlet and collected in a beaker at the outlet. Meanwhile, a pressure gauge (MA059, MEASUREMAN) and an air pressure regulator (850-AC, ControlAir Inc.) were used to control the air pressure from a compressed air source directed at the control channel. The pressure regulator was used to adjust the control pressure and the liquid collected in the outlet beaker was massed after a known collection time (i.e., 10 s) on a digital analytical balance (XS204, Mettler-Toledo) to determine the flow rate. The air pressure in the control channel was increased in ∼3 kPa increments until the outlet flow rate neared 0 µL s^-1^, indicating that the valve was closed.

### ELISA-chip nitrocellulose assay preparation

The assay was designed based on lateral flow nitrocellulose membranes (Vivid 120, no. VIV1202503R; Pall Corporation, Port Washington, USA) that were cut to 3 × 12 mm^2^ (width × length) with a pointed base using a film cutter (Cameo 3, Silhouette Portrait, Lindon, USA). The membranes were then spotted using an inkjet spotter (sciFLEXARRAYER SX, Scienion) with a 2.5 × 1 mm^2^ (width × length) test and control line spaced 5 mm apart. The test line was spotted with 100 µg mL^-1^ of anti-human IFN-γ antibody in a 0.22-µm filtered 1X PBS buffer by programming the release of 350 pL droplets in a 25 × 4 line array; spotting was done over 40 passes, wherein each pass covered alternating positions on the line array to allow for spots to dry between passes. Similarly, the control line was spotted with 50 µg mL^-1^ of BSA-biotin with 8 passes covering alternating positions for each pass. The spotted membranes were dried at 37°C for 1 h, then blocked in a solution of 0.1% Tween 20 in 1X PBS supplemented with 5% BSA by dipping and wetting the membranes in a tray containing excess blocking buffer placed on an orbital shaker at 100 rpm for 1 h at room temperature. The blocked membranes were left to dry at 37°C for 1 h, followed by overnight storage in an air-tight container with desiccant at 4°C, then used within 48 h of protein spotting. The nitrocellulose assay strip was mounted onto the chip and sandwiched between 4 absorbent pads (Electrophoresis and Blotting Paper, Grade 320, Ahlstrom-Munksjo Chromatography) laser cut to be 10 × 2.4 mm^2^ (width × length). The drainage channel of the chip was mounted with a 1.5 × 3.5 mm^2^ (width × length) glass fiber (G041 SureWick, Millipore Sigma), then both the glass fiber and nitrocellulose strips were clamped into place using a custom 3D printed compressive clip.

### Assay protocol

The assay solutions for IFN-γ detection were prepared in a wash and diluent buffer of 0.05% Tween 20 in 1X PBS supplemented with 5% BSA. Recombinant human IFN-γ protein was spiked in the buffer at concentrations of 0, 10^0^, 10^1^, 10^2^, 10^3^, 10^4^, 10^5^, and 10^6^ pg mL^-1^, followed by preparing reagent solutions including anti-human IFN-gamma biotinylated antibody at 1 µg mL^-1^ and streptavidin-poly-horseradish peroxidase (pHRP) at 25 µg mL^-1^. To prepare the assay substrate solution, SIGMAFAST™ DAB (3,3’-diaminobenzidine) tablets were dissolved in 5 mL Milli-Q water, then 0.22-µm filtered prior to running the assay.

### Videos and image processing

3D images of the microfluidic devices were obtained by micro-computed tomography (µCT) (SkyScan 1172, Bruker, Kontich, Belgium) at a pixel size of 8 μm. Images were reconstructed using CT Analyzer (CTAn v.1.18, Bruker, Kontich, Belgium) and orthogonal projections were visualized and measured in ImageJ2 Ver. 2.9.0/1.53t (public domain software, National Institute of Health, Bethesda, Maryland, United States). 3D microscopy images were taken with a stereomicroscope (SteREO Discovery.V20, Zeiss, Baden-Württemberg, Germany). Videos and images were recorded to characterize flow using dyed water on either a Panasonic Lumix DMC-GH3K or Sony α7R III camera. Assay membranes were imaged using a flatbed scanner (Epson Perfection V600) with the SilverFast 8 software at 600 dpi in a 48-bit RGB format, then imported to ImageJ2 for 16-bit grayscale colorimetric line intensity readouts. The readouts were normalized to rescale the colorimetric intensity from 0-65535 gray values to 0-1 relative signal intensities.

## Results and discussion

**Figure 1** shows the process flow including a low-cost LCD 3D printer, a custom PEGDA-based ink (PLInk) optimized for LCD 3D printing, some microfluidic devices fabricated in this study, and the device design that closes the rapid prototyping cycle.

**Figure 1.**
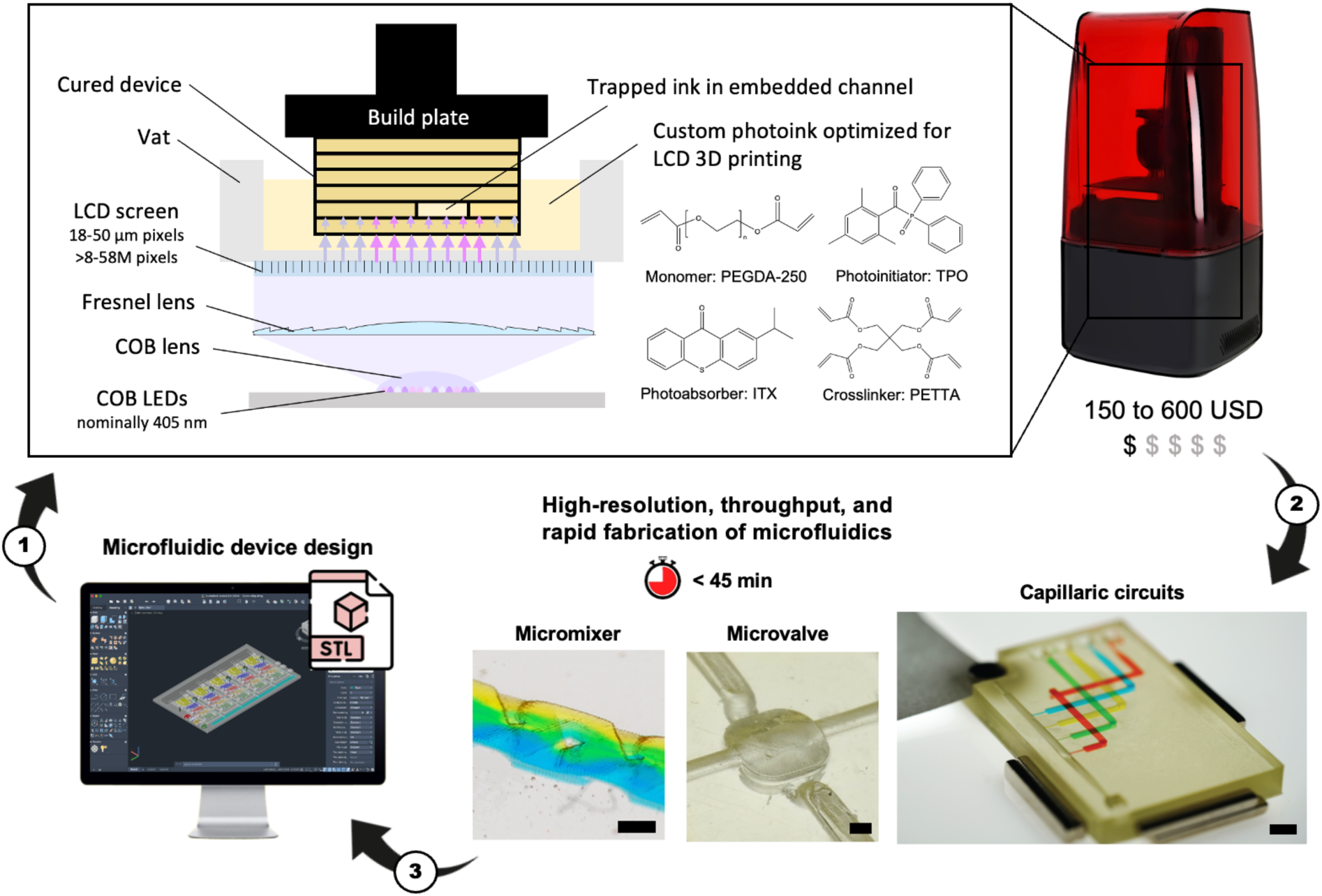
Low-cost LCD 3D printing of microfluidic devices and the rapid prototyping cycle. The workflow including (1) manufacturing on low-cost LCD 3D printers using a custom PEGDA-based ink (PLInk) optimized for LCD 3D printing, (2) directly manufactured microfluidic devices that can be tested and characterized, and (3) inform design improvement for the next iteration. Pixel sizes of 18-50 µm and print areas of up to 218 × 122 mm^2^ afford high print resolution over large areas. Photoinks optimized for LCD 3D printing with reduced sensitivity to light heterogeneity and low viscosity enable the direct manufacture of microfluidic chips including open and embedded microchannels with a lateral resolution <100 µm and vertical features as thin as 22 µm in <45 min. Scale bars = 500 µm.

### Design of ink for LCD 3D printing

Embedded microfluidic channels are designed as narrow voids in a block of solid ink. To create these voids, the design of an ink formulation consists of monomers, light-responsive additives (i.e., photoinitiator to catalyze the reaction, photoabsorbers to absorb excess energy), and crosslinkers. Polymerization must proceed efficiently layer-by-layer, i.e., within the defined thickness of each layer while both avoiding under-polymerization of the current layer, and over-polymerization of uncured ink in voids of the preceding layers.

The design of PLInk was based on our prior ink formulations for 385 nm DLP 3D printing,^7, 16^ but adapted for LCD-based photopolymerization by considering the light heterogeneity, low irradiance, and 405 nm illumination wavelength. Based on our prior inks, PEGDA-250 was selected as the monomer due to its low viscosity, low protein adsorption, inherent cytocompatibility, and compatibility with solvents such isopropyl alcohol for efficient removal of uncured ink in embedded microchannels. Diphenyl(2,4,6-trimethylbenzoyl)phosphine oxide (TPO) was selected again as the photoinitiator due to its low cytotoxicity and an activation peak between 380-425 nm, as well as 2-isopropylthioxanthone (ITX) as the photoabsorber due to its broad absorbance peak between 350-425 nm, and known optical transparency, unlike other photoabsorbers such 2-nitrophenyl phenyl sulfide (NPS), Sudan-1, or UV absorbing dyes with poor cytocompatibility and yellow-orange tints. Due to the low irradiance of LCD 3D printers, we added pentaerythritol tetraacrylate (PETTA) crosslinker to increase reactivity (discussed further below). Each of these ink components individually met suitability for a 405 nm illumination source, **Figure 2a**.

**Figure 2.**
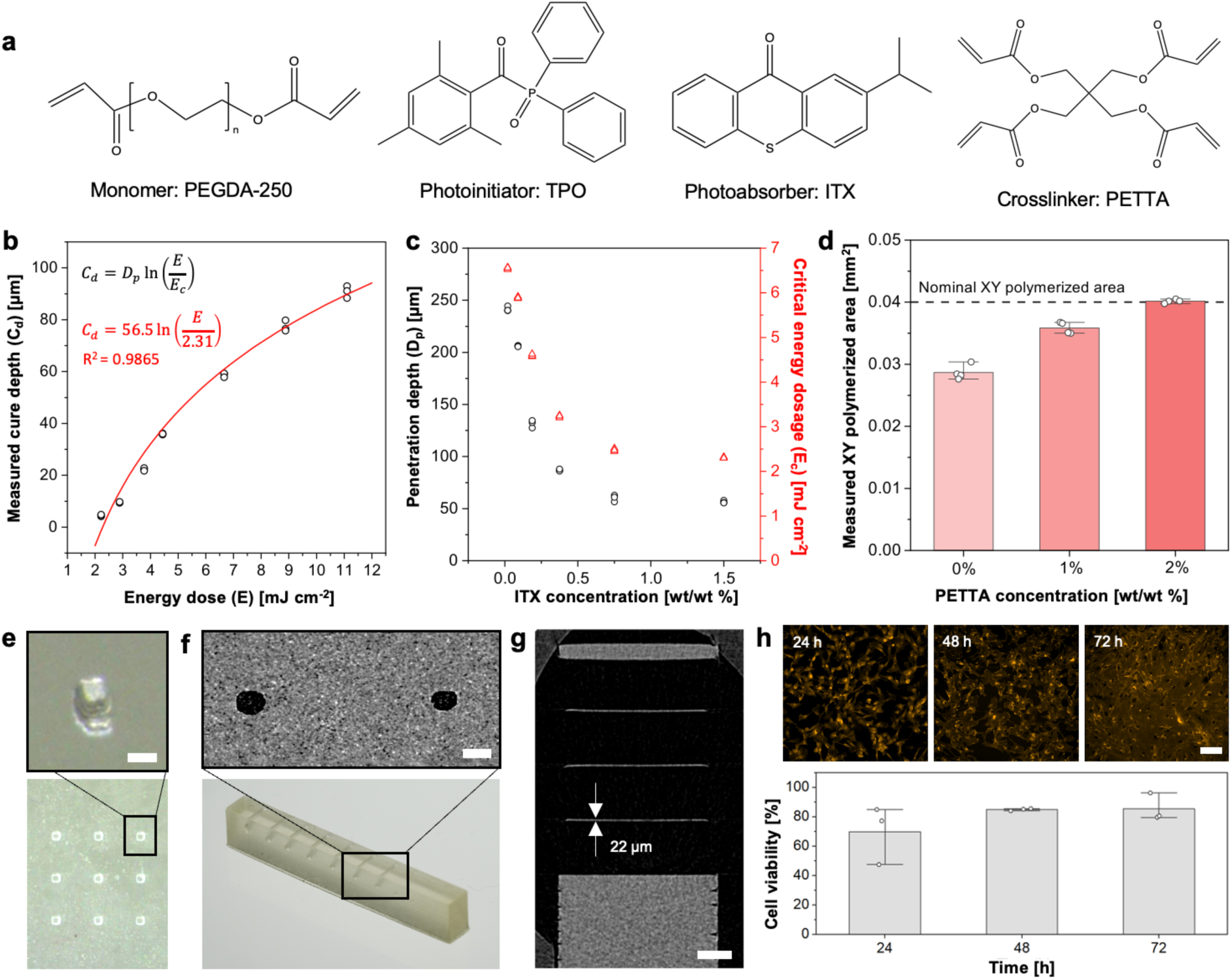
PLInk characterization for microfluidic LCD 3D printing. (a) Formulation of PLInk containing PEGDA-250, TPO, ITX and PETTA. (b) Jacob’s working curve showing the cure depth as a function of the energy dose for 1.5% ITX and yielding *D_p_* = 56.5 µm and *E_c_* = 2.31 mJ cm^-2^. (c) *D_p_* and *E_c_* values derived as in (b) for different concentrations of ITX showing a plateau for ITX > 0.75%. (d) Pillar printed area compared to nominal area of 0.2 × 0.2 mm^2^ as function of crosslinker concentration. Data shows mean ± standard deviation (STD) of four replicates. (e) 3D printed pillars showing features as small as ∼3 × 3 pixels (nominal printer pixel size = 28.5 × 28.5 µm^2^). Scale bar = 100 µm. (f) Monolithic circular microchannels with a corresponding µCT scan with channel cross-sections radii of ∼125 µm (left) and ∼110 µm (right) (nominal printer pixel size = 35 × 35 µm^2^). Scale bar = 250 µm. (g) µCT image of a 22-µm thick embedded membrane (nominal printer pixel size = 18 × 18 µm^2^). Scale bar = 500 µm. (h) Cell viability at different time points of HUVECs expressing actin-mCherry co-cultured with 3D printed PLInk rings. Data shows mean ± STD of three biological replicates. Scale bar = 100 µm.

We confirmed photocuring by Fourier Transform Infrared-Attenuated Total Reflectance (FTIR-ATR) spectroscopy on uncured and 405 nm cured PLInk samples. A broader peak at 1200 cm^-1^ was observed for the cured ink, consistent with carbon-carbon bond formation between adjacent PEGDA and PETTA acrylate groups, Supplementary **Figure S1**.

Next, to mitigate the effects of light inhomogeneity, we sought to characterize the photopolymerization of the ink as function of total energy dosage and varying ITX photoabsorber concentration from 0 to 1.5%; the latter being the maximal concentration at which ITX could readily be dissolved. The fabrication of embedded microchannels, i.e., voids, is predicated on precise control and understanding of the (measured) cure depth, *C_d_*, to both avoid cross-linking of uncured ink trapped inside the microchannel while ensuring curing of the working layer. *C_d_* is experimentally measurable and varies as function of the total energy, *E,* according to Jacob’s working curve:^12, 22, 29, 30^

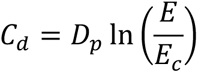

Where *D_p_* is the penetration depth of light at which the light intensity of incident light is reduced by a factor 1/*e*,^22^ and *E_c_* is the critical energy dosage corresponding to the minimum required energy to initiate photopolymerization. *E* is simply *t_e_*, the exposure time multiplied by *P*, the irradiance:

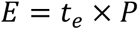

The LCD 3D printer irradiance was measured to be 2.23 mW cm^-2^. The thickness of polymerized ink for 1.5% ITX as function of energy dosage was fitted with Jacob’s curve to derive both *D_p_* = 56.5 µm and *E_c_* = 2.31 mJ cm^-2^, **Figure 2b**. The same experiment was repeated for varying concentrations of ITX, and the resulting *D_p_* and *E_c_* derived, **Figure 2c**. As expected, *D_p_* decreased with increasing ITX concentration. Interestingly, *E_c_* also decreased with increasing photoabsorber, suggesting that ITX contributes not only to light adsorption, but also to more effective photopolymerization of the ink. We also observed that the slope was flattened for higher ITX concentration, meaning that the variation in the thickness of photopolymerized ink as a result of light illumination inhomogeneity would be minimized. Hence, high ITX concentrations were optimal for LCD 3D printing. While we observed a lower plateau in both *D_p_* and *E_c_* once ITX concentrations reached 0.75%, we chose 1.5% as the optimal condition to minimize susceptibility to variations in ITX concentration. The UV-Vis absorbance spectra showed that reducing the penetration depth improved the ink efficiency by increasing the absorbance near ∼405 nm, thus matching the illumination wavelength of the 3D printer, Supplementary **Figure S2**.

To balance precision, material sensitivity and print speed, and while considering printer pixel size, we set the print layer thickness (and model slicing) to 20 µm. This satisfied the requirement for printing embedded microchannels of slice layer thickness = 0.3-1 × *D_p_* formulated by Nordin and colleagues.^22^ PLInk also allowed for rapid photopolymerization with an exposure time *t_e_* of 1.3-1.8 s for a *C_d_* of ∼20 µm. As an example of the benefits of lower *D_p_*, for a change in energy dosage of 7-9 mJ cm^-2^, the layer thickness variation with 1.5% and 0.02% ITX would be ∼20 µm and ∼80 µm, respectively, Supplementary **Figure S3**. Commercial inks typically favor a high *D_p_* (>179 µm),^31^ which has the advantage of printing thicker layer slices and faster print times, but are inadequate for printing embedded microchannels and susceptible to variable cured thickness with a non-uniform light source.

To improve printing fidelity, we supplemented the PLInk formulation with PETTA with four additional acrylate groups to increase the availability of polymerizable groups and speed up the formation of an interconnected polymer network. We empirically adjusted the PETTA concentration by measuring the printed area of 0.2 × 0.2 mm^2^ pillars with a 28.5-μm pixel size LCD 3D printer. Incomplete photopolymerization was visualized by tracking underfilling of the nominal pillar shape and by the distortion and bending of the pillars.^32^ The PETTA concentration was increased until the nominal XY pillar area matched the 3D printed design, which was achieved at a value of 2%, **Figure 2d**. Pillar printing confirmed suitable mechanical stability of the print without collapse and good dimensional accuracy, as illustrated with an array of ∼3 × 3 pillars, **Figure 2e**.

### PLInk performance characterization

PLInk is based on PEGDA-250 with a comparatively low viscosity of ∼16 mPa s, thus making it suitable for microchannel fabrication. Indeed, following photoexposure of a layer, the retraction of a relatively flat print attached to the vat bottom will create suction force; next, ink needs to flow into the growing gap, and immediately flow out of the gap as the print is lowered back onto the vat bottom to expose the next layer, all of which would benefit from a low viscosity ink. Notably, despite its high viscosity (∼700 mPa s at 25°C),^33^ the addition of PETTA at low concentrations did not impact the native viscosity of PEGDA-250, and thus vastly outperformed commercial inks (∼200-500 mPa s) in this respect, Supplementary **Table S2**. The low viscosity also facilitates printing of fine features as it reduced the risks of mechanical failure caused by suction and adhesion to the vat bottom. Coupling low viscosity and low *D_p_*, the cured PLInk was smooth with a surface roughness of ∼500 nm, which suggests favourability for intricate microchannel fabrication, Supplementary **Section S1** and **Figure S4**.

To assess suitability for microfluidic device fabrication, we evaluated the resolution of the designed PLInk formulation by printing open channels with decreasing size and were able to print features as small as ∼75 × 75 µm^2^ with a 35-μm pixel size LCD 3D printer, Supplementary **Figure S5**. We performed µCT scans of the device to evaluate the printing accuracy; we measured the printed open channel size and found it to be within 2.8% of the nominal dimension.

To assess our ability to 3D print embedded microchannels, we similarly evaluated the printing of progressively smaller rectangular and circular channels that were embedded a depth at least ten times greater than the *D_p_*. Embedded rectangular channels down to ∼170 × 220 µm^2^ (width × height) were printed using a 35-µm pixel size LCD 3D printer, Supplementary **Figure S6**. The smallest rectangular embedded conduits were within 2.7% of their nominal size. We found that high aspect ratio (height [H] / width [W] > 1) channels were limited by the pixel resolution of the LCD screen, i.e., typically 3-4 pixels because of scattering, non-parallel illumination, and possible photoinitiator diffusion.^34, 35^ Meanwhile, the height of low aspect ratio microchannels (H/W < 1) was limited by the optical penetration (the shortest embedded channel ∼2.3 × *D_p_*).^12^ Circular conduits are notably of interest to minimize capillary edge flow (also called filaments),^7^ and embedded conduits with circular cross-section and radius as small as ∼110 µm was printed with a dimensional accuracy within 1.5% of their nominal dimension, **Figure 2f** and Supplementary **Figure S7**. A shallow *D_p_* also benefits the printing of thin embedded membranes due to fine control over the cured thickness and a sharp transition between cured and uncured layers. Vertical embedded channels designed with a series of ever thinner membranes were 3D printed down to a thickness as low as ∼22 µm within a single exposure to demonstrate free-standing membrane fabrication, **Figure 2g** and Supplementary **Figure S8**.

Further, we evaluated the cytocompatibility of the ink by co-culturing 3D printed PLInk with human umbilical vein endothelial cells (HUVECs) according to the ISO 10993-5:2009 standard for implantable medical devices. A primary cell line was selected due to specific but rigorous culturing conditions for cells with high sensitivity to their environment and a limited passage number. We 3D printed 8 × 3 mm^2^ (diameter × thickness) rings and thoroughly washed any unreacted photoactive elements (details in the Methods), then co-incubated the PLInk rings with cells in a single well with shared media for 72 h.^15^ After 72 h, we found >80% cell viability, meeting the threshold for a cytocompatible material and demonstrating suitability for cell culture microfluidic device fabrication, **Figure 2h**.

In summary, the optimized and low viscosity PLInk formulation for LCD 3D printing was found to be suitable for high-resolution and dimensionally accurate printing of smooth structures including posts, open and embedded microchannels, (embedded) membranes, and to be cytocompatible, making it amenable for a broad range of applications, and notably in microfluidics as explored below.

### LCD 3D printing of embedded microfluidic mixer

We re-designed a recently published micromixer implementing the Baker’s transformation and made by direct laser micromachining of two separate parts for direct LCD 3D printing.^36^ In the original design, the micromixer was assembled from two substrate devices with open conduits made by direct laser machining, chemical wet etching, and bonding, thus forming a closed, interconnected weaving flow path. For LCD 3D printing, the micromixer was designed as a single digital model of an embedded micromixer including (i) overhanging wedges to split the fluidic streams horizontally and progressively, and (ii) an embedded pillar to create an interface before vertically recombining the two fluidic streams, **Figure 3a**. The micromixer was designed with 310 × 310 µm^2^ cross-sections that interweaved, merged into a large conduit of 900 × 900 µm^2^, and then split again, and so on. The shaped pillars measured 203 µm (≅ 6 pixels, nominal 3D printer pixel size = 35 × 35 µm^2^) along its longest dimension and 306 µm (≅ 9 pixels) along its widest dimension, **Figure 3b**. The pillar extended across the full height of the mixer (45 layers of 20 µm each), which necessitated a sufficiently high energy dosage to fully crosslink the pillar as well as overhanging, suspended structures and preserve their integrity during the build plate movements, while at the same time preventing photopolymerization of PLInk in the embedded weaving conduits. µCT scans of the 3D printed device confirmed the fidelity and integrity of the pillar and the overhangs, and of internal corners and sharp edges that split, guide, and merge the fluidic streams, **Figure 3c(i)**. A finite element method numerical simulation that solved the steady-state concentration field of two fluidic streams illustrated the importance of the microarchitecture to horizontally split and vertically recombine the flows for cross-sections along the length of the mixing unit, **Figure 3c(ii)**.

**Figure 3.**
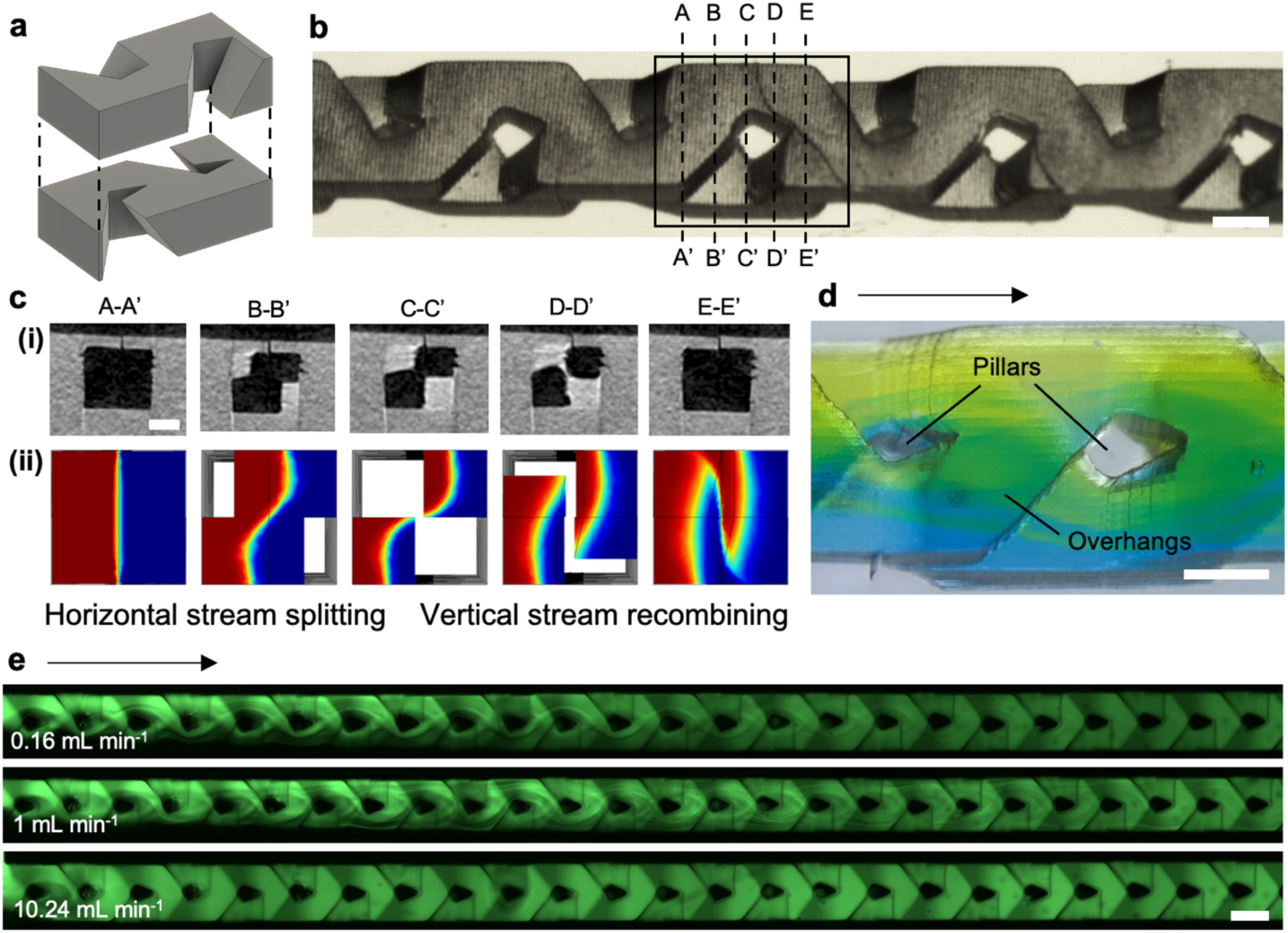
Embedded microfluidic mixer. (a) Schematic representation of two fluidic channels combined into a single mixing unit. (b) Stereomicroscope image of four 3D printed mixing units showing overhangs and pillar formation in an embedded device. Scale bar = 500 µm. (c) µCT cross-sections of the microchannel with a corresponding numerical simulation showing the mixing principle based on horizontal stream splitting and vertical stream recombining. Scale bar = 300 µm. (d) Stereomicroscope image of a single mixing unit showing yellow and blue fluidic streams split into ever thinning striations by the microarchitecture. Arrow shows the direction of flow. Scale bar = 500 µm. (e) Mixing of 10 μM fluorescein with clear water at a flow rate ≅ 0.1, 1, and 10 mL min^-1^, corresponding to a Reynold’s number ≅ 1.85, 18.5, and 185, respectively. Arrow shows the direction of flow. Scale bar = 500 µm.

Owing to the transparency of the device, mixing could be visually tracked through the entire height of the channel. Using water with yellow and blue dyes allowed for visual tracking of the mixing and the observation of striations as the streams folded and recombined within the micromixer, **Figure 3d**. We further assessed the mixing performance with a fluorescent dye (10 µM fluorescein) in one of the streams and tracked the fluorescence intensity along the length of the mixer by fluorescence microscopy. The progression from two separate streams to complete mixing was visible from the intensity profile that progressed from a step function to a flat, homogeneous distribution of the dye across the width of the micromixer, **Figure 3e** and Supplementary **Figure S9**. When investigated over a range of laminar flow rates (0.01-10 mL min^-1^), we observed the efficiency of mixing decreased with increasing flow rates, as expected because the time for diffusive mixing decreases. Interestingly, we observed that for flow rates >1 mL min^-1^ the mixing efficiency did not decrease, but instead improved again, which we attribute to inertial effects and recirculation. The mixing performance was concordant with the laser-manufactured mixer and the Baker’s transformation principle.^36, 37^ Across three replicate devices, we quantified the mixing efficiency to be 92-99%, confirming the successful printing and operation of the 3D printed device, Supplementary **Figure S10**, **Section S2**. The micromixer illustrates the potential of LCD 3D printing for producing complex embedded structures that are not easily manufactured by more traditional micromachining methods.

### LCD 3D printing of an embedded membrane microvalve

Next, we 3D printed an embedded membrane microvalve. Elastomeric microvalves made of polydimethylsiloxane (PDMS) and manufactured by replica molding have been widely used and adopted for microfluidics.^38, 39^ Recently, direct manufacturing of embedded free-standing membranes by 3D printing has been demonstrated using DLP 3D printers.^29, 40^ We LCD 3D printed an embedded membrane with a valve seat modelled based on existing ones comprising a 40-µm-thick membrane with a diameter of 1.7 mm and ∼100 µm above a 500-µm-wide ridged valve seat. An embedded control channel overlaid orthogonally above the membrane and the valve seat in the flow channel was used for membrane actuation by pressurization, **Figure 4a-b**. All the channels were 3D printed with a unique inlet and outlet to facilitate precursor ink removal and avoid post-processing fabrication steps. The valve seat in the form of a thin curved ridge improved printability compared to a solid ‘bowl’ shape that might lead to incomplete ink removal while providing reliable valve closure upon actuation. µCT images of the valve revealed a fully released, free-standing membrane, **Figure 4c**, Supplementary **Video S1**. The measured thickness on the µCT images (with 8 µm pixel resolution) was ∼43 µm, closely matching the design.

**Figure 4.**
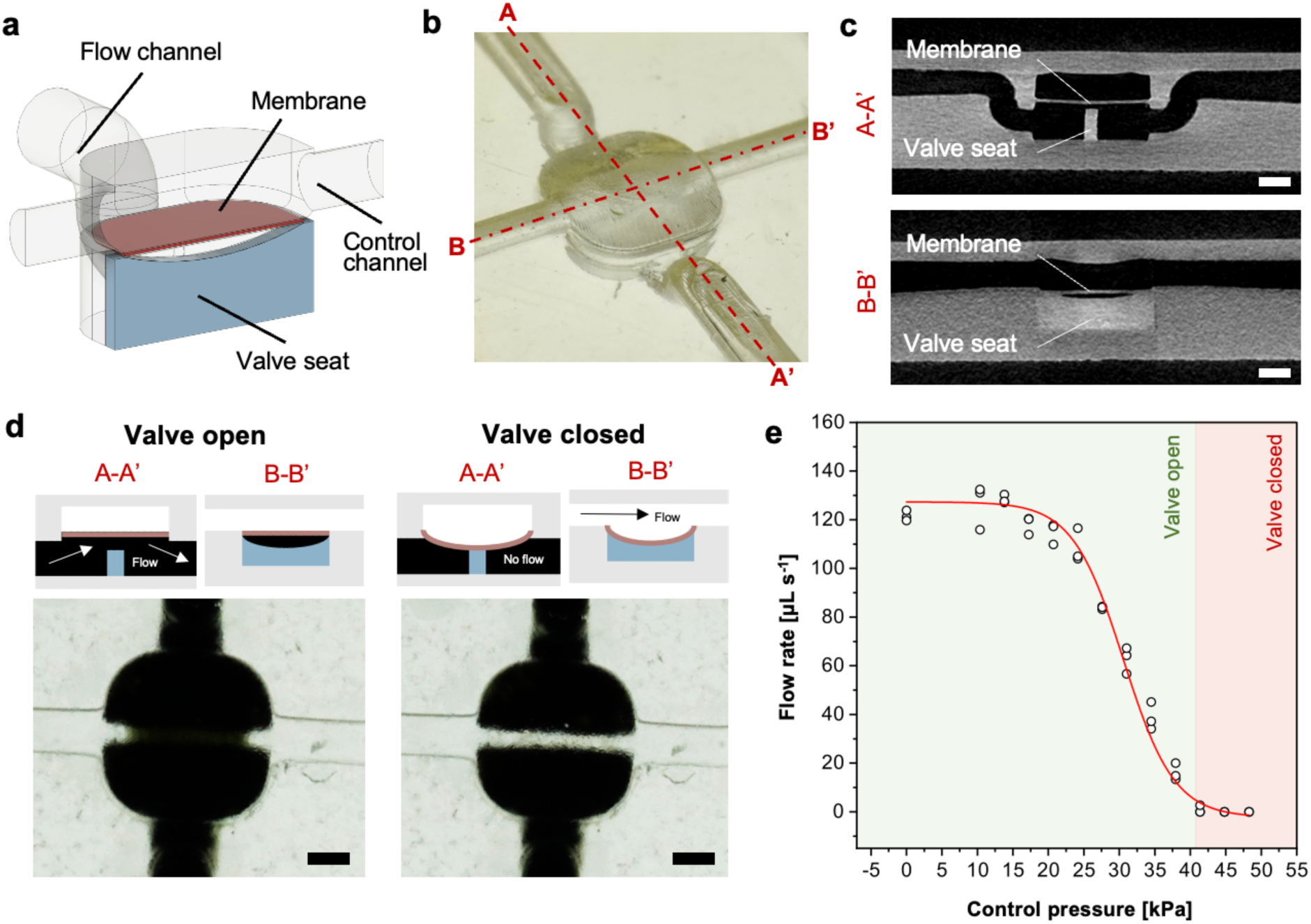
Embedded membrane microvalve. (a) Microvalve schematic and (b) photograph showing flow channel, control channel, a ∼43 µm thick membrane, ridged valve seat forming a separation wall with a 100 µm gap to the membrane at the centre (nominal 3D printer pixel size = 28.5 × 28.5 µm^2^). Actuation of the membrane by pressurization in the control channel leads to deflection onto the valve seat and closure of the flow channel. (c) Orthogonal µCT views of the 3D printed membrane and valve seat. Scale bars = 500 µm. (d) Top view images of the open and closed valve with schematics showing cross-sections of the valve according to the labels in (b). With an open valve shown on the right, the black water flows through the channel, while for a closed valve, the pneumatically deflected membrane is sealed onto the valve seat and stops black water flow. Scale bars = 500 µm. (e) Flow rate in the flow channel as a function of the pressure in the control channel showing the gradual closing of the valve and flow stop at ∼41 kPa. Data points are measurement collected from three different devices. Line is a guide to the eye.

Water spiked with a black dye was flown through the microvalve to visually assess whether the valve was open (i.e., flow channel junction was visually black) or closed (i.e., junction visually clear). The valve was designed to be open at rest, and as the compressed air pressure was increased in the control channel, the membrane deflected to form a seal with the valve seat, interrupting the flow of the black water, **Figure 4d**.

The mechanical properties of 3D printed PLInk were assessed by tensile testing yielding a Young’s modulus of 68 ± 3 MPa, Supplementary **Figure S11**. Compared to elastomeric membranes, PLInk’s Young’s modulus was ∼10× higher than PDMS; therefore, a thin (∼40-50 µm), 1.7 mm diameter membrane was predicted to deflect ∼100 µm at a control pressure of ∼45 kPa to seal the valve, Supplementary **Section S3**. The control pressure was increased incrementally while the flow was monitored and flow stop observed at ∼41 kPa, **Figure 4e**. The experimental valve closing pressure was thus in good agreement with the prediction, and the variation could be attributed to imprecision in the gap between the membrane and the valve, in the thickness of the membrane, or incomplete curing of the membrane that might make it more pliable. Overall, both the reproducibility of the closing pressure across all valve replicates, and the agreement to theory were consistent. While we did not assess the durability under cyclical stress loading, the durability of 3D printed membranes based on low molecular weight PEGDA inks was demonstrated by Folch and colleagues,^40^ suggesting that the PLInk membrane will also be suitable, or could be made suitable, for cyclical loading. These results indicate that LCD 3D printing can be used for making thin, compliant, and mechanically actuated embedded elements such as membrane microvalves.

### LCD 3D printing of an ELISA-on-a-chip capillaric circuit – an ELISA-chip

CCs operate by structurally encoding fluidic operations using capillary valves for fluidic operation and capillary flow for self-filling, and function thanks to a controlled, moderate hydrophilicity. We previously developed hydrophilic inks for DLP 3D printing of functional CCs with embedded channels and with contact angles with water ∼35° owing to the use of hydrophilic acrylic or methacrylic acid additives.^7^ The contact angle with water of PLInk was ∼65-70°, which while being moderately hydrophilic, was too high for reliable capillary self-filling, Supplementary **Figure S12a**. The photopolymeriziation of acrylic or methacrylic acid groups competes with crosslinking by PEGDA acrylate groups, and thus requires higher light energy doses, which would lead to much higher exposure times for low irradiance LCD 3D printers. Previously, plasma activation had also been used, but depends on access to a plasma chamber, and only provides temporarily hydrophilicity for select materials.^18, 19^ Hence, instead of increasing the surface energy of the microchannels, we opted to reduce the surface tension of the aqueous solutions by adding surfactants (i.e., Tween 20) and thereby reducing the contact angle to as low as ∼46° with 0.05% Tween 20 and ∼31° with 0.1% Tween 20, thus meeting the requirements for CC operation, Supplementary **Figure S12b-c**. Considering that the use of surfactants in immunoassays is common to reduce non-specific binding, their addition to the solutions does not compromise the suitability of CCs for typical biological applications.

To illustrate the reliability of LCD 3D printing, we designed a CC with a microfluidic chain reaction (MCR)^18^ implementing an ELISA-on-a-chip akin to the ones made previously using DLP 3D printing of open microchannels followed by sealing with a hydrophobic pressure adhesive transparent cover.^18, 19^ The ELISA-chip was developed for a new target, with adjusted geometries for LCD 3D printing, and importantly with a reduced time-to-result while maintaining high sensitivity, **Figure 5a**. The target was interferon (IFN)-γ, a cytokine critical to the immune response against a wide range of infections,^41^ and which is notably used in the IFN-γ release assays as a biomarker for tuberculosis infection.^42, 43^ The microfluidic assay was based on a classical ELISA sandwich immunoassay using a capture antibody, a biotinylated detection antibody, and a streptavidin-enzyme conjugate (poly-horseradish peroxidase, pHRP). While in conventional well-plate ELISAs soluble substrates are used, for on-chip applications with a nitrocellulose membrane and under active flow conditions, precipitating substrates are required for localized accumulation of the enzymatically oxidized substrate, such as 3,3′-diaminobenzidine tetrahydrochloride (DAB), in the presence of pHRP and hydrogen peroxide, **Figure 5b**. A nitrocellulose membrane spotted with an anti-IFN-γ capture antibody was connected to the ELISA-chip that encoded an 8-step assay for automated, sequential flow of wash buffers and reagents. As in the DLP 3D printed ELISA-chip design, functions for on-chip aliquoting were integrated to facilitate the operations for untrained users, **Figure 5c**.

**Figure 5.**
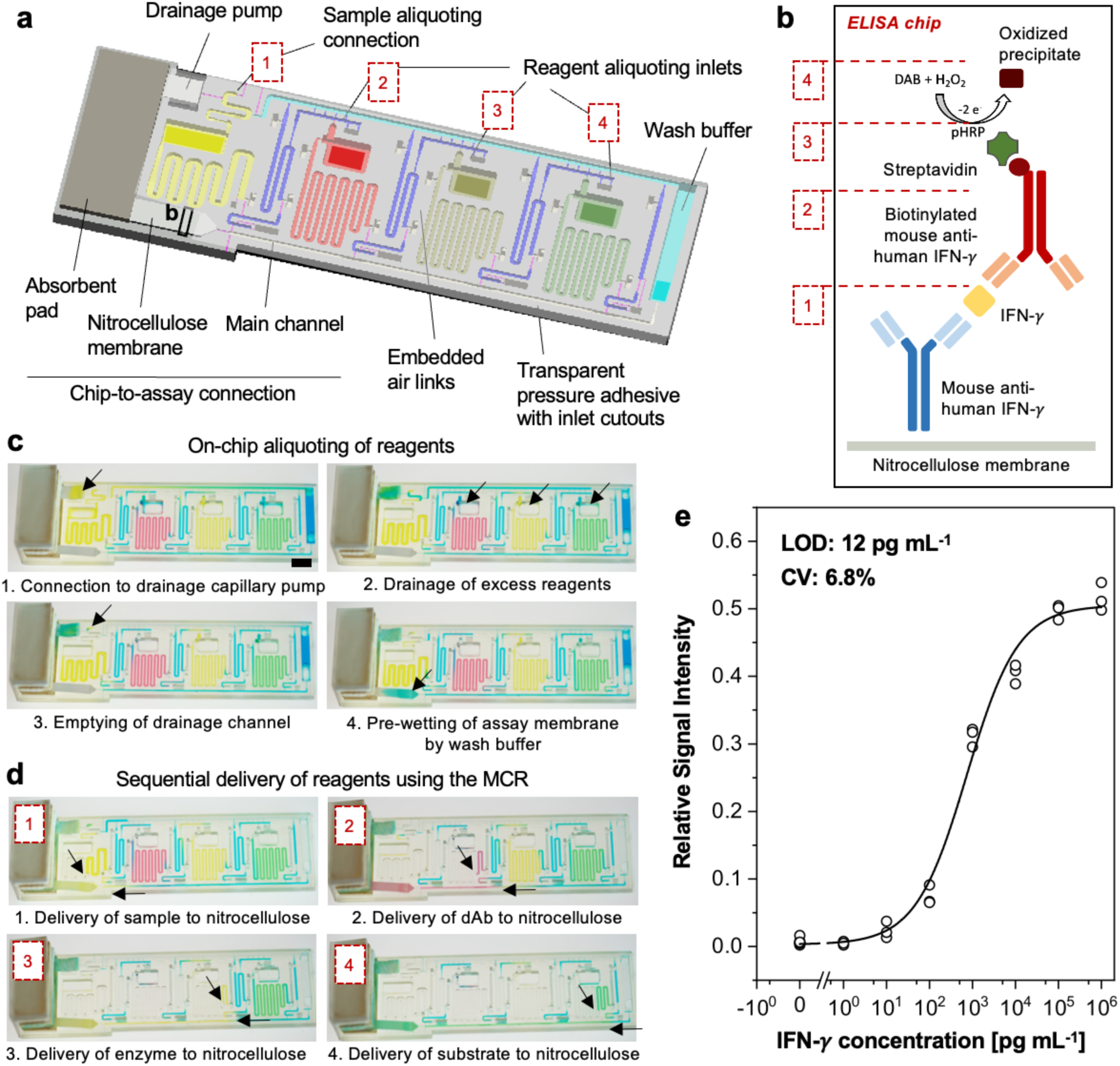
LCD 3D-printed microfluidic ELISA chip. (a) ELISA-on-a-chip designed for LCD 3D printing with structurally encoded sequential delivery of reagents to autonomously perform an assay and coupled with a built-in chip-to-assay connection. (b) ELISA workflow showing the sandwich immunoassay designed for the detection of IFN-γ by sequentially delivering assay reagents and wash buffer to a nitrocellulose membrane pre-spotted with anti-human IFN-γ capture antibody; (1) sample containing IFN-γ, (2) biotinylated anti-human IFN-γ detection antibody, (3) streptavidin-conjugated enzyme pHRP, and (4) enzyme substrate in the presence of hydrogen peroxide to generate the colorimetric readout. (c) Autonomous CC workflow for on-chip reagent aliquoting by metering the correct reagent volumes and drainage of the excess, followed by (d) MCR-based sequential delivery of assay reagents with wash steps in between. Arrows show the direction of flow. Scale bar = 5 mm. (e) Binding curve of the on-chip assay for the detection of IFN-γ with a limit of detection of 12 pg mL^-1^ (CV: 6.8%) across triplicate chips for each tested concentration point; line shows a 4-point logistic fit.

The lower limit of detection of the previous ELISA-chip^19^ outperformed rapid tests (e.g., lateral flow assays), but the assay time was longer at 1 h 15 min. Thus, we sought to reduce the assay time for the LCD 3D printed ELISA-chip. The incubation times were structurally encoded by the volume of reagents that flowed over the test zone (see discussion on assay optimization below for further details), the capillary pressure of the pump (i.e., absorbent pad and glass fiber conjugate pad backing the nitrocellulose membrane), and the flow resistance of the functional connections that linked each reservoir to the main channel. The capillary pressure coming from an absorbent pad backing the nitrocellulose membrane was the same as a single pump was used to wick all the reagents. Compared to our previous ELISA-chip design that also had a glass fiber conjugate pad mounted the nitrocellulose and served both as a fluidic connection to the chip and an immediate capillary pump to wick reagents over the nitrocellulose, the glass fiber was considered a source of analyte loss due to protein adsorption over the assay run time. To remedy these limitations, we connected the nitrocellulose membrane to the ELISA-chip directly. Without the glass fiber, the chip-to-assay connection was re-designed as a gradual opening with a weak stop valve designed to break when the liquid front arrived at the end of the channel and wetted the nitrocellulose membrane; pre-wetting with buffer bridged the ELISA-chip’s liquid interface with the absorbent pad, and facilitated a connection to the capillary pump that subsequently began to wick the reagents over the nitrocellulose assay test zone. Finally, to adjust the flow rate, we increased the functional connection cross-sections to 200 × 200 µm^2^ across the entire chip. These changes reduced reagent loss and provided a suitable flow speed for consistent fluidic performance, which afforded the option to reduce reagent volumes and the time-to-result down to 48 min, **Figure 5d**, Supplementary **Video S2**.

We evaluated the flow reproducibility of the new LCD 3D printed ELISA-chip by timing each of the sequential steps in three replicate chips, **Table 1**. The flow of sample, which contains only limited concentration of analyte is the most critical step when considering assay reproducibility and LOD, and the one that necessitated high reproducibility. Other steps, such as detection antibody and enzyme are provided in excess concentration and hence variation of flow time is not expected to significantly affect the assay result. Likewise, precise incubation time for wash steps are not as critical as long as reagents are flowed and flushed across the nitrocellulose membrane. The comparatively high variability for the DAB incubation time could arise as a result of the precipitate formed on the test strip, especially at higher concentrations of IFN-γ, which could affect the flow properties of the strip.

**Table 1.** LCD 3D printed ELISA-chip steps, reagent, volume and timing of automated assay.

| | Reagent | Vol. [$\mu\text{L}$] | Time ± STD [s] |
| --- | --- | --- | --- |
| 1 | Sample (IFN-γ) | 75 | $875 \pm 16$ (CV: 1.9%) |
| 2 | Wash buffer, PBST 0.05% + 5% BSA | 15 | $152 \pm 21$ (CV: 14.2%) |
| 3 | Biotinylated detection antibody | 45 | $513 \pm 21$ (CV: 3.9%) |
| 4 | Wash buffer | 15 | $144 \pm 19$ (CV: 12.8%) |
| 5 | Streptavidin-pHRP | 45 | $518 \pm 34$ (CV: 6.5%) |
| 6 | Wash buffer | 15 | $146 \pm 17$ (CV: 17.5%) |
| 7 | Enzyme substrate DAB | 45 | $504 \pm 52$ (CV: 10.3%) |
| 8 | Wash buffer | 5 | $79 \pm 25$ (CV: 31.6%) |

The assay portion of the ELISA-chip was optimized using a design of experiments approach, which enabled the optimization of multiple assay parameters simultaneously since the optimal concentration of one parameter would dictate the optimal of another in a classical sandwich immunoassay, and served to establish the relative contribution of each parameter.^44^ We evaluated a capture antibody spotting concentration of 50, 100 and 200 µg mL^-1^ and both a detection antibody and pHRP concentration of 1, 5, and 25 µg mL^-1^ at a fixed sample concentration of 100 ng mL^-1^. Using the Taguchi method for design of experiments,^45^ the selection led to nine experiments to determine significantly impacting assay factors, Supplementary **Table S3**. From the results, we evaluated the significance of each factor using analysis of variance and found that the capture antibody concentration was a significant parameter (*p* < 0.05) for the assay performance, and the weighted contribution of the capture antibody concentration was found to be 47%, which was higher than the other factors, i.e., detection antibody (25%), and pHRP (24%), Supplementary **Table S4**. Altogether, this indicated that a critical point in reducing assay time while preserving the sensitivity was to increase the capture antibody spotting density. To that end, we kept the reagent volumes relatively low, i.e., sample volume was 75 µL which took ∼14 mins to flow, and ensured that all the reagent were being delivered to the nitrocellulose membrane with no loses on a connecting glass fiber; meanwhile, we increased the spotting density of capture antibody from our original ELISA-chip by nearly 10-fold, resulting in 0.7 µL of 100 µg mL^-1^ capture antibody spotted on a thin 3 × 1 mm^2^ (width × length) line on the nitrocellulose membrane. Taking the relative contribution of each parameter into consideration, the optimal IFN-γ assay thus required flowing the detection antibody at 1 µg mL^-1^ and pHRP at 25 µg mL^-1^ over a fixed incubation time encoded by 45 µL of both reagents. The assay was optimized with minimal wash steps to reduce the assay run time and served to prevent pre-mixing of reagents in the main channel so they would only conjugate at the test zone.

Following both optimization of the fluidic performance and the nitrocellulose assay, we evaluated the ELISA-chip over a wide concentration range of IFN-γ and achieved a limit of detection as low as 12 pg mL^-1^. With a 6.8% CV, our LCD 3D printed ELISA-chip showed consistent performance, **Figure 5e**, Supplementary **Figure S13**. These results indicate the suitability of low-cost LCD 3D printing for the fabrication of ready-to-use CC chips that automate complete assays with lab-grade accuracy and short time-to-result.

### Digital manufacturing by LCD 3D printing

The ELISA-chips previously fabricated by DLP 3D printing were fabricated one chip at a time, but following the redesign to decrease the footprint, and the much larger print bed of the 4K LCD 3D printer (>10M pixels over 143 × 89 mm^2^), 5 ELISA-chips could be printed at once in <45 min. We optimized LCD 3D printings settings for high feature density manufacturing, focusing on layer rest times, retraction/descent speeds, and build plate rise between layers. While open designs have limited contact points to the vat bottom (e.g., bucky ball structures, scaffold designs, triply periodic minimal surfaces, etc.), 3D printing of large flat surfaces create significant challenges during the retraction of the print from the flexible vat bottom due to a strong suction force that could lead to mechanical damage of intricate features. Slower retraction speeds, and longer rest times provided sufficient time to replenish the vat for relatively flat ELISA-chips, while quick descents were used to minimize the print time. A low-rise height of 1-2 mm between retraction and descent prevented bubble formation, while further minimizing print times. These optimizations improved print resolution for high feature density and prevented common failures such as 3D printing of distinct adjacent channels as joint. Using PLInk, and based on the cost of research-grade ingredients, the ELISA-chip would a cost of ∼2 USD per device, Supplementary **Table S5**. The low capital cost and low material cost enable affordable fabrication of autonomous ELISA-chip devices globally, especially in low- and middle-income countries with limited access to traditional manufacturing and a high incidence of infectious disease.

Using optimized settings and CCs with a smaller footprint, many more chips can be printed at once. To illustrate this possibility, we designed a CC with a footprint of 34 × 22 mm^2^ and capillary retention burst valves with increasing burst-thresholds encoded by decreasing cross-sections (down to ∼180 × 180 µm^2^, nominal 3D printer pixel size = 28.5 × 28.5 µm^2^) for the sequential delivery of 4 solutions via embedded and weaving conduits,^46^ **Figure S14**, Supplementary **Video S3**. 42 visually accurate CC systems could be digitally manufactured by 3D printing in a single 45-min run. This corresponds to a manufacturing throughput of 448 chips per 8 h per 3D printer, Supplementary **Figure S15**. Hence using ten 8K LCD 3D printers at a total capital cost of ∼6000 USD, and that could be operated by one technician, ∼13,440 functional CCs could be printed per 24 h, immediately upon receipt of the digital design file. Hence LCD 3D printing offers the potential for on-demand, low capital cost, and high throughput manufacturing.

## Conclusion

We presented the use of low-cost photopolymerization LCD 3D printing for the fabrication of microfluidic devices using PLInk, optimized for rapid polymerization under low irradiance, 405 nm illumination, and reliable printing of embedded microchannels (and thin membranes) despite illumination inhomogeneity. The effect of ITX photoabsorber concentration on ink photopolymerization and notably *D_p_* and *E_c_* was characterized for 20-µm-layer-by-layer printing. Posts with lateral resolution of 75 µm, embedded membranes 22 µm thin, and embedded microchannels with rectangular cross-sections of 170 × 220 µm^2^ and round cross-sections with 110 µm radius were 3D printed. Further, we demonstrated that microfluidic devices previously made by other methods, such as laser machining, replica molding and DLP 3D printing, can now be fabricated using LCD 3D printing, including an embedded micromixer, a membrane microvalve, and an ELISA-chip for IFN-γ detection. Finally, we showcased the potential for throughput manufacturing by leveraging the large build area to digitally manufacture >13,000 CCs in 24 h using 10 3D printers.

Future work could explore the concurrent variation of TPO photoinitiator, ITX photoabsorber, and PETTA crosslinker concentration to better understand their interplay with regards to *D_p_*, *E_c_,* and printing accuracy, and choose the optimal mixture based on specific applications and criteria. Photoabsorbers with a higher absorption efficiency than ITX at 405 nm could also be explored; however, ITX offers excellent transparency and higher wavelength absorption often comes at the expense of optical transparency and yellow-orange tinted devices. Finally, mapping the light heterogeneity of the LCD 3D printers by the end user, and the tools to do that, would open the door to digital correction of the illumination heterogeneity by programming the 3D printer, and further improve the resolution achievable both with commercial and custom photoinks.

We may expect that 3D stereolithography printer manufacturers driven by market pressure will continue to increase pixel numbers and concomitantly reduce pixel size, all while preserving the affordability of LCD 3D printers, which will further increase their appeal and adoption. We foresee that some of the greatest opportunities lie in improving the photoinks for LCD (and more generally stereolithography) 3D printing, which are in their infancy. While here we showed the application of LCD 3D printing to microfluidics that were primarily designed based on prior manufacturing technologies, opportunities arise to re-design and ideate microfluidic systems that leverage the strength of LCD and stereolithography 3D printing.

Digital manufacturing by LCD 3D printing is as simple as downloading a file and printing it, thus circumventing the need for specialized machinery and advanced training, while enabling customizability and rapid design iterations by the end user. The advent of low-cost and easy-to-use 3D printers compared to traditional manufacturing methods enables the fabrication of open and embedded microscopic features by anyone, anywhere, thus democratizing access to high-resolution fabrication and reducing the entry barrier for many potential users. The combination of low-cost, high-resolution 3D printers, and readily 3D printable designs enable the realization of low-cost and distributed digital manufacturing.

## Acknowledgements

We acknowledge Yongjun Xiao for assistance with µCT at the Centre for Bone and Periodontal Research, Mohsen Ketabi and Gwénaël Chamoulaud for assistance in profilometry, viscometry, and FTIR-ATR at the NanoQAM Research Center, and Lucie Riffard for assistance in tensile testing at the McGill Structures and Composite Materials Laboratory. H. S. acknowledges a CGS-M NSERC, an FRQNT master’s research scholarship and a McGill BME recruitment award. V. K. acknowledges an FRQNT doctoral research scholarship. G. K. acknowledges an FRQNT doctoral research scholarship and a McGill BME recruitment award. Y. M. acknowledges a McGill BME recruitment award. A. S.-K. acknowledges FRQNT postdoctoral fellowship. D. J. acknowledges support from a Canada Research Chair in Bioengineering.

## Author contributions

V. K. designed the ink formulation, V. K. and H. S. characterized the ink and assessed printability for LCD 3D printing. M. L. S. performed the biocompatibility assay. H. S., G. K., and Y. M. printed and characterized the micromixer. G. K. and H. S. printed and characterized the microvalve. A. S.-K. contributed to the embedded sequential delivery chip. H. S. designed and optimized the ELISA-chips. H. S. prepared the figures and analyzed the data. H. S., V. K., and D. J. prepared the initial draft of the manuscript, with feedback from all the authors. H. S. and D. J. edited and reviewed the manuscript drafts. D. J. provided funding, and conceptualized, supervised, and administered the project.

## Competing interests

The authors have no conflicts of interest to declare.

## Data availability

3D design files are uploaded to Thingiverse and Printables (https://www.thingiverse.com/junckerlab/collections and https://www.printables.com/@JunckerLab_743461).

Data not presented in the article or supplementary material will be available upon request. Correspondence and requests for materials should be addressed to: David Juncker 6500-740 Dr Penfield Avenue Montreal, QC. CA. H3A 0G1 Phone: +1 (514) 398 7676, Fax: +1 (514) 398 1790

## Notes

### Competing Interest Statement

The authors have declared no competing interest.

