## Supplementary Information for "High-resolution low-cost LCD 3D printing of microfluidics"

**Supplementary calculations**

**Section S1.** Measurement of surface roughness

**Section S2.** Calculation of mixing index

**Section S3.** Calculation of the predicted valve closing pressure

**Supplementary tables**

**Table S1.** Summary of 3D printing settings

**Table S2.** Viscosity of 3D printing inks

**Table S3.** Summary of Taguchi Design of Experiments assay results

**Table S4.** Analysis of variance and percent contribution from Taguchi Design of Experiments

**Table S5.** CC-chips cost breakdown using PLInk

**Supplementary figures**

**Figure S1.** FTIR-ATR spectra

**Figure S2.** UV-Vis spectra

**Figure S3.** Cure depth as a function of the energy dose

**Figure S4.** Surface roughness of 3D-printed parts using PLInk

**Figure S5.** Open channel 3D printing

**Figure S6.** Rectangular channel 3D printing

**Figure S7.** Circular channel 3D printing

**Figure S8.** *In situ* membrane 3D printing

**Figure S9.** Fluorescent images of microfluidic mixing at different volumetric flow rates
**Figure S10.** Mixing indices
**Figure S11.** Tensile testing of 3D-printed parts using PLInk
**Figure S12.** Contact angle
**Figure S13.** Nitrocellulose membranes from LCD 3D printed ELISA-chip for IFN-γ detection
**Figure S14.** Monolithic CC
**Figure S15.** Throughput manufacturing of CCs

**Supplementary videos**

**Video S1.** μCT scan of microvalve
**Video S3.** LCD 3D printed ELISA-chip
**Video S3.** Embedded CC device

### Supplementary calculations

#### Section S1. Measurement of surface roughness

The surface roughness of a 3D printed part was measured using profilometry according to the ISO 4287 and 4288 standards. Sample prints that were 25 × 75 × 2 $mm^3$ (width × length × height) were assessed for surface roughness. The mean surface roughness is given by the average distance between the mean line and all the data points over a minimum sampling length of 0.8 mm; the surface roughness ($R_a$) is calculated as follows:

$$R_a = \frac{1}{n}\sum_{i=1}^{n}|y_i| \qquad \text{(Equation S1)}$$

where $y$ denotes all the sampled points, and $n$ denotes the number of sampled points that are iterated over an increment $i$. Over three unique sampling lengths, the average surface roughness of PLInk on the low-cost 3D printers was 530 ± 110 nm (**Figure S4**).

#### Section S2. Calculation of mixing index

The mixing efficiency was assessed by measuring the fluorescent intensity of a pre-mixed solution versus a solution mixed in the microchannels. The captured images of the mixed solutions were analyzed in ImageJ by taking the fluorescent intensity readout over the length of the channel outlet. For all the flow rates and the given channel geometry, we calculated the corresponding relative mixing index (RMI) as a metric for efficiency as follows:

$$RMI = \left[1 - \frac{\sigma}{\sigma_0}\right] \times 100\% = \left[1 - \frac{\sqrt{\frac{\sum_{i=1}^{n}\left(x_i - x_{avg}\right)^2}{n}}}{\sqrt{\frac{\sum_{i=1}^{n}\left(x_{0i} - x_{avg}\right)^2}{n}}}\right] \times 100\% \qquad \text{(Equation S2)}$$

where $\sigma$ and $\sigma_0$ are the STDs of the fluorescent intensities, $x_i$ is the normalized intensity at point $i$ at the outlet, $x_{0i}$ is the normalized intensity at point $i$ at the inlet, $x_{avg}$ is the normalized intensity for a completely mixed solution, and $n$ denotes the number of points sampled. With this metric, RMI = 100% represents a state with complete mixing while RMI = 0% represents a state with no mixing.

#### Section S3. Calculation of the predicted valve closing pressure

Based on known valve dimensions and known material properties, we can estimate the pressure required to deflect the membrane and make contact with the valve seat to subsequently close the valve. For the valve we present here, the predicted valve closing pressure ($P$) can be calculated as follows:[1]

$$P_{closing} = \left(\frac{5.33}{1-v^2}\left(\frac{y}{t}\right) + \frac{2.6}{1-v^2}\left(\frac{y}{t}\right)^3\right)\left(\frac{Et^4}{r^4}\right) \qquad \text{(Equation S3)}$$

where *E* is the Young's modulus (~68 MPa), *v* is the Poisson ratio (~0.9), *r* is the membrane radius (~1.05 mm), *t* is the membrane thickness (~43 μm), and *y* is the gap height that the membrane must deflect (~100 μm). This yields a pressure of ~45 kPa to close the valve, which is in good agreement with the physical experiment.

**Supplementary tables**

**Table S1.** Summary of 3D printing settings.

| | **Calibration devices** (Figure S5-7) | **Calibration devices** (Figure S8) | **Microfluidic mixer** (Figure 3) | **Membrane microvalve** (Figure 4) | **ELISA-chip** (Figure 5) | **Sequential delivery chip** (Figure S14-15) |
|---|---|---|---|---|---|---|
| Printer (All from Elegoo) | Mars 3 Pro 4K | Mars 4 Ultra 9K | Mars 3 Pro 4K | Saturn 2 8K | Mars 3 Pro 4K | Saturn 2 8K |
| Light engine* | 36 COB LED + Fresnel lens | COB LED + Fresnel lens | 36 COB LED + Fresnel lens | 48 COB LED + Fresnel lens | 36 COB LED + Fresnel lens | 48 COB LED + Fresnel lens |
| Light uniformity and angle* | 92% | 92%, <5º | 92% | Not reported | 92% | Not reported |
| Pixel size [μm$^2$]* | 35 × 35 | 18 × 18 | 35 × 35 | 28.5 × 28.5 | 35 × 35 | 28.5 × 28.5 |
| Pixel number [pixels]* | 4098 × 2560 ≅ 10M | 8520 × 4320 ≅ 36M | 4098 × 2560 ≅ 10M | 7680 × 4320 ≅ 33M | 4098 × 2560 ≅ 10M | 7680 × 4320 ≅ 33M |
| Layer height [mm] | 0.020 | 0.020 | 0.020 | 0.020 | 0.020 | 0.020 |
| Exposure time [s] | 1.3 | 1.3 | 1.6 | 1.5 | 1.8 | 1.3 |
| Bottom exposure time [s] | 7 | 7 | 10 | 10 | 7 | 10 |
| Rest time [s] | 3 | 3 | 3 | 3 | 5 | 3 |
| Lifting speed [mm min$^{-1}$] | 50 | 50 | 30 | 65 | 50 | 50 |
| Retract speed [mm min$^{-1}$] | 210 | 210 | 60 | 80 | 210 | 210 |
| Anti-aliasing | Yes | Yes | Yes | Yes | No | No |
| Total print time [min] | ~51 | ~51 | ~39 | ~31 | ~42 | ~40 |

Chip-on-board LEDs are called COB. * indicates manufacturer information taken from their website at https://www.elegoo.com.

**Table S2.** Viscosity of 3D printing inks

| **Ink** | **Viscosity [mPa s]*** |
|---|---|
| PLInk containing 0% PETTA crosslinker | 16.2 |
| PLInk containing 0.5% PETTA crosslinker | 15.8 |
| PLInk containing 1% PETTA crosslinker | 15.0 |
| PLInk containing 2% PETTA crosslinker | 15.6 |
| *Anycubic upgraded ABS-like clear resin* | *224* |
| *Monocure 3D crystal clear resin* | *551* |
| *Luvantix 3DMaterials clear resin* | *528* |

* viscosity was measured at 21ºC

**Table S3.** Summary of Taguchi Design of Experiments assay results

| | **Capture antibody [µg mL$^{-1}$]** | **Detection antibody [µg mL$^{-1}$]** | **pHRP [µg mL$^{-1}$]** | **Relative signal intensity [0-1]** |
|---|---|---|---|---|
| 1 | 50 | 1 | 1 | 0.42 |
| 2 | 50 | 5 | 5 | 0.59 |
| 3 | 50 | 25 | 25 | 0.63 |
| 4 | 100 | 1 | 5 | 0.55 |
| 5 | 100 | 5 | 25 | 0.63 |
| 6 | 100 | 25 | 1 | 0.51 |
| 7 | 200 | 1 | 25 | 0.61 |
| 8 | 200 | 5 | 1 | 0.59 |
| 9 | 200 | 25 | 5 | 0.67 |

**Table S4.** Analysis of variance and percent contribution from Taguchi Design of Experiments

| Source | Degree of freedom | Sum of squares | F ratio | p-value | Contribution |
|---|---|---|---|---|---|
| pHRP | 2 | 114.3 | 18.9 | 0.050 | 25% |
| dAb | 2 | 104.5 | 12.4 | 0.054 | 24% |
| cAb | 2 | 200.0 | 33.3 | 0.029* | 47% |

* for significance level $\alpha$=0.05

See Luo *et al.*[2] for more details.

**Table S5.** CC-chips cost breakdown using PLInk

| **Item** | **Vendor** | **Price [USD]** | **Quantity** | **Cost per chip [USD]** | |
|---|---|---|---|---|---|
| | | | | **Embedded CC device (20 × 30 × 3 mm$^3$ ≅ 1.7 mL)** | **ELISA-chip (87 × 27 × 2.8 mm$^3$ ≅ 5 mL)** |
| PEGDA-250 | Sigma-Aldrich | \$142 | 500 mL | \$0.72 | \$2.15 |
| ITX | Fisher Scientific | \$205 | 25 g | \$0.01 | \$0.03 |
| TPO | Sigma-Aldrich | \$159 | 50 g | \$0.01 | \$0.01 |
| PETTA | Sigma-Aldrich | \$110 | 250 mL | \$0.02 | \$0.04 |
| | | | Total | \$0.76 | \$2.23 |

**Supplementary figures**

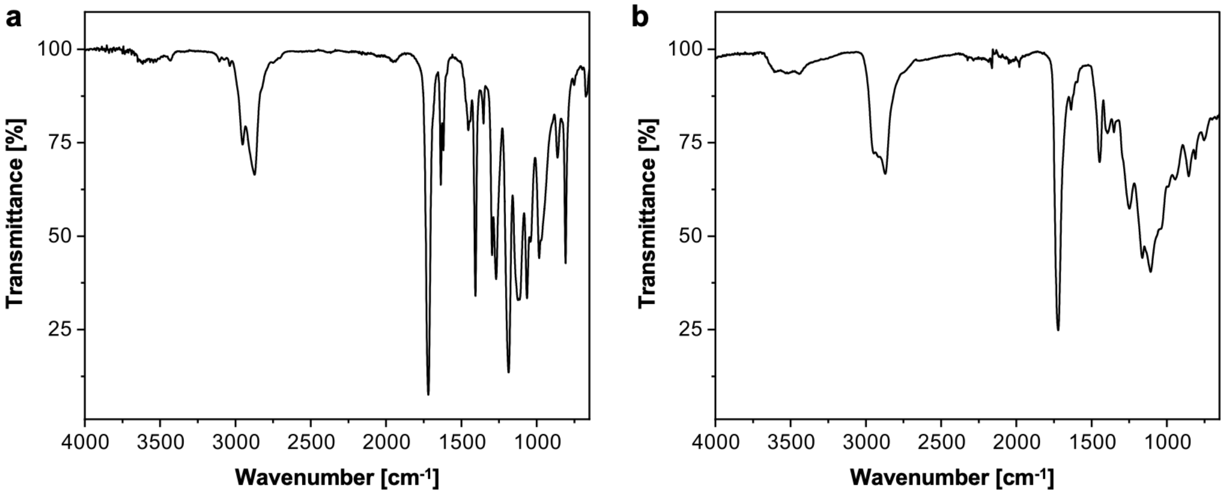


**Figure S1. FTIR-ATR spectra.** Functional group characterization using FTIR-ATR of the (a) uncured precursor PLInk and (b) cured PLInk following 405 nm UV exposure.

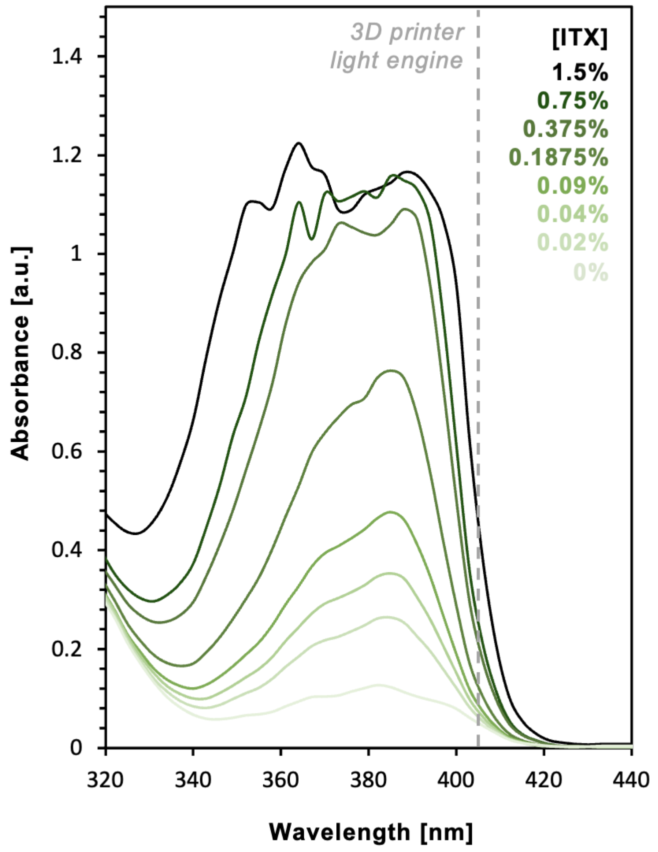


**Figure S2. UV-Vis spectra.** Higher ITX concentration increased the absorbance at the illumination wavelength of 3D printer (i.e., 405 nm) to improve the efficiency of PLInk for LCD 3D printing.

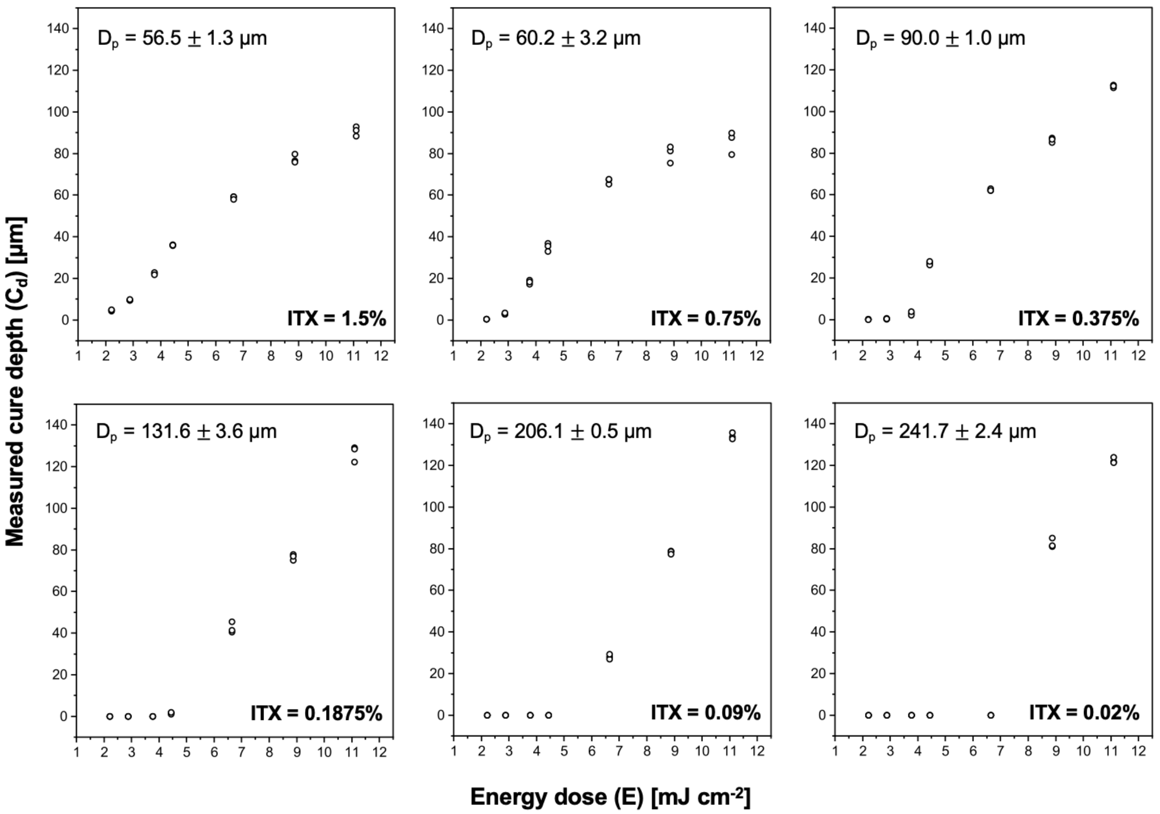


**Figure S3. Cure depth as a function of the energy dose**, for different concentrations of ITX to determine the penetration depth of light for PLInk. Data shows mean ± STD across three replicates and was acquired using the Elegoo Mars 3 Pro 3D printer.

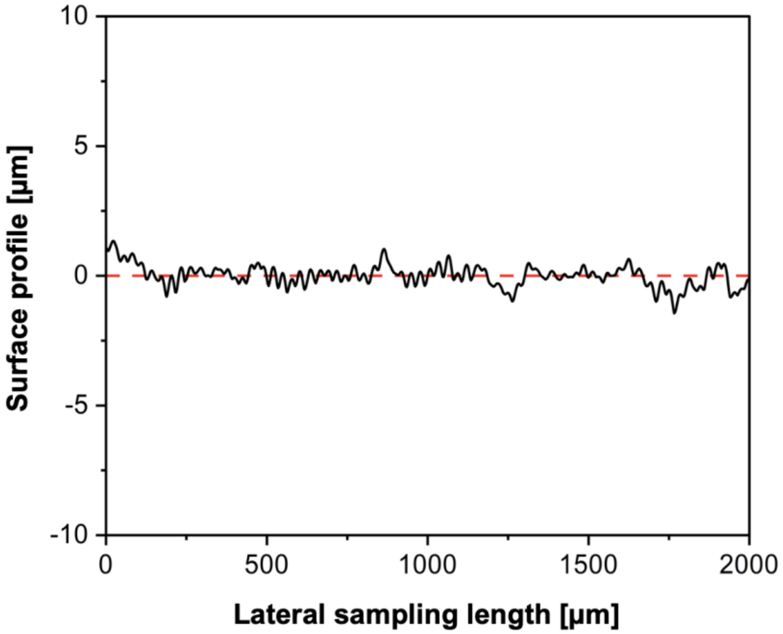


**Figure S4. Surface roughness of 3D printed parts using PLInk.** Surface topography given along a 12.5 µm line width using a stylus profilometer to calculate a mean surface roughness of 530 ± 110 nm across three replicates. Data was acquired using the Elegoo Mars 3 Pro 3D printer.

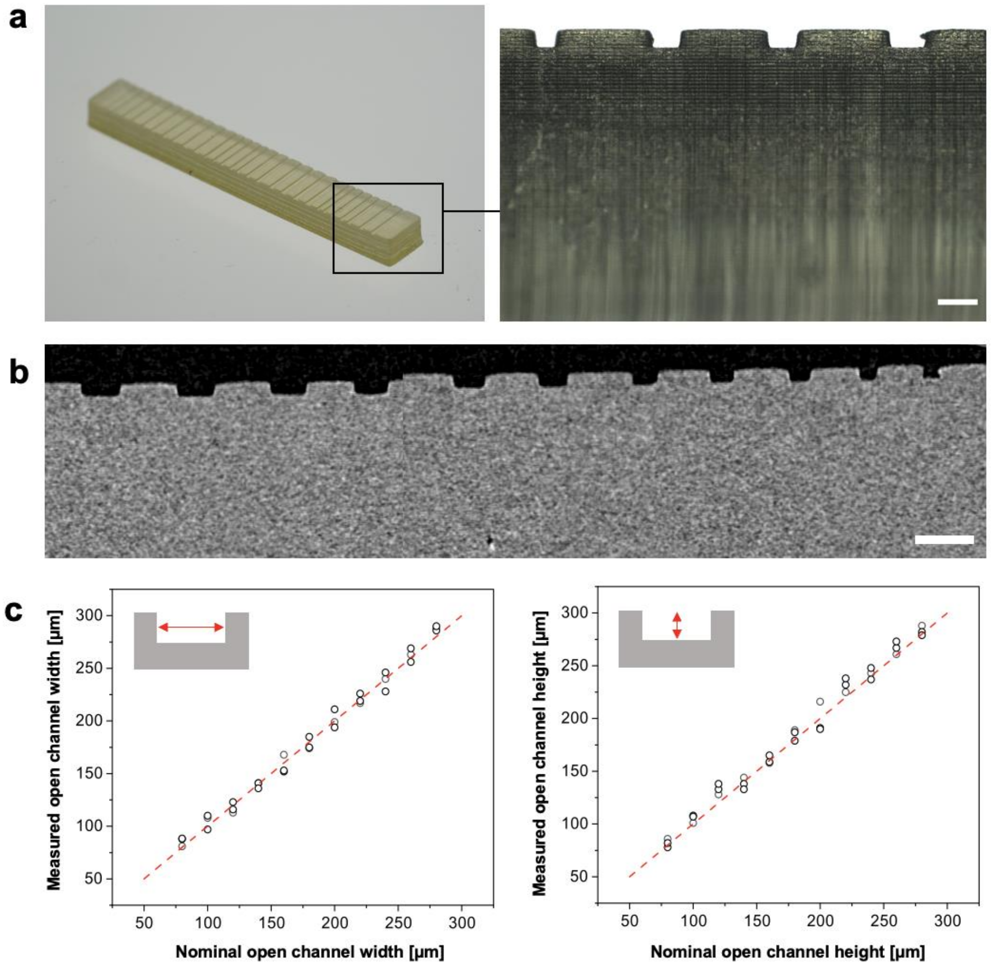


**Figure S5. Open channel 3D printing.** (a) Open channel calibrator device showing varying channel dimensions to assess the resolution, inset shows ~75 × 75 µm$^2$ open channel at the surface of the device. Scale bar = 200 µm. (b) µCT side view of the open channel calibrator device. Scale bar = 500 µm. (c) Dimensional accuracy between design file and printed channel widths and depths across three unique devices. Dashed line shows the case where the nominal value = measured value.

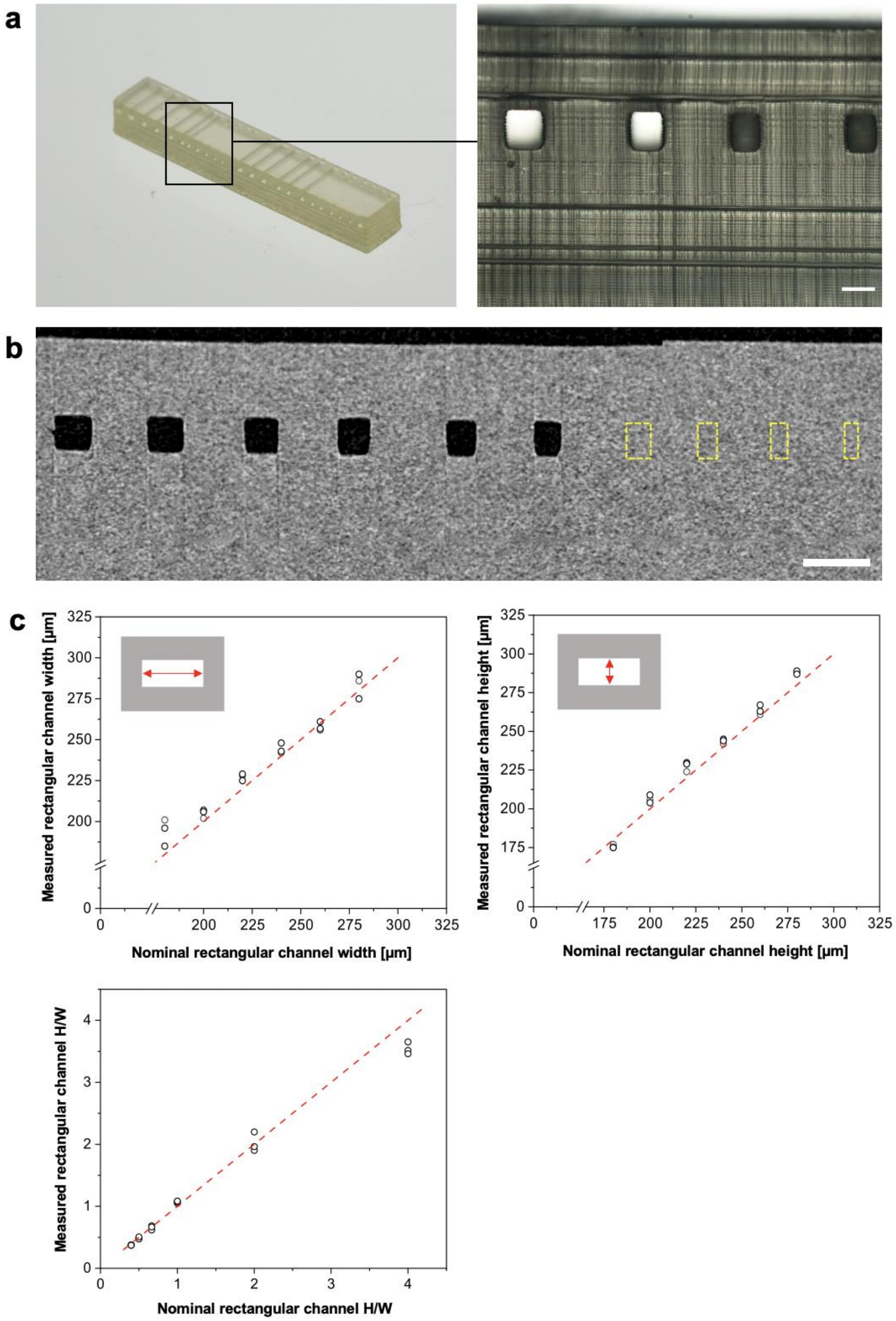


**Figure S6. Rectangular channel 3D printing.** (a) Rectangular channel calibrator device showing varying channel dimensions to assess the resolution, inset shows ~170 × 220 µm$^2$ rectangular channel embedded into the device. Scale bar = 200 µm. (b) µCT side view of the rectangular channel calibrator device showing clogged channels in yellow dashed outlines. Scale bar = 500 µm. (c) Dimensional accuracy between design file and printed channel widths and depths across three unique devices; aspect ratio dimensional accuracy is given as the height (H) divided by width (W) of rectangular microchannels. Dashed line shows the case where the nominal value = measured value.

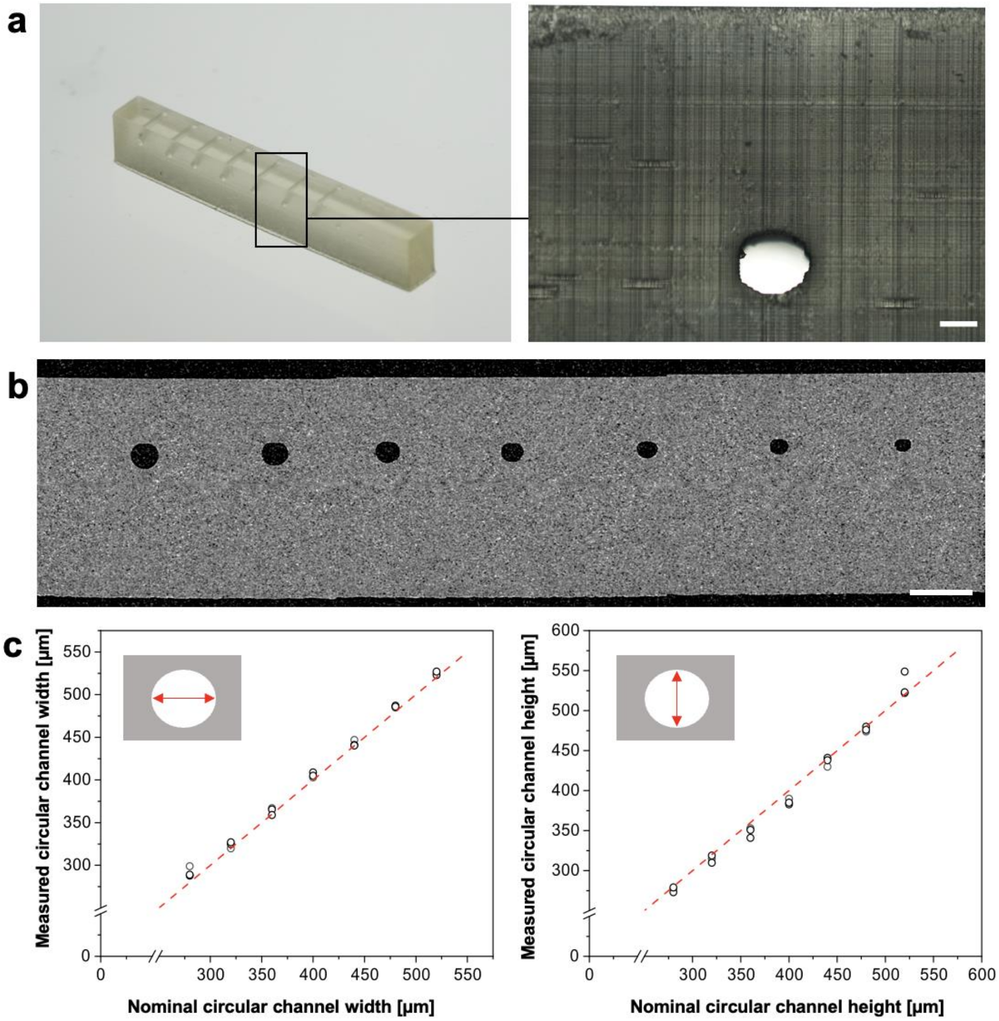


**Figure S7. Circular channel 3D printing.** (a) Circular channel calibrator device showing varying channel diameters to assess the resolution, inset shows a ~125 µm radius embedded channel. Scale bar = 200 µm. (b) µCT side view of circular channel calibrator device. Scale bar = 1 mm. (c) Dimensional accuracy between design file and printed channel widths and depths across three unique devices. Dashed line shows the case where the nominal value = measured value.

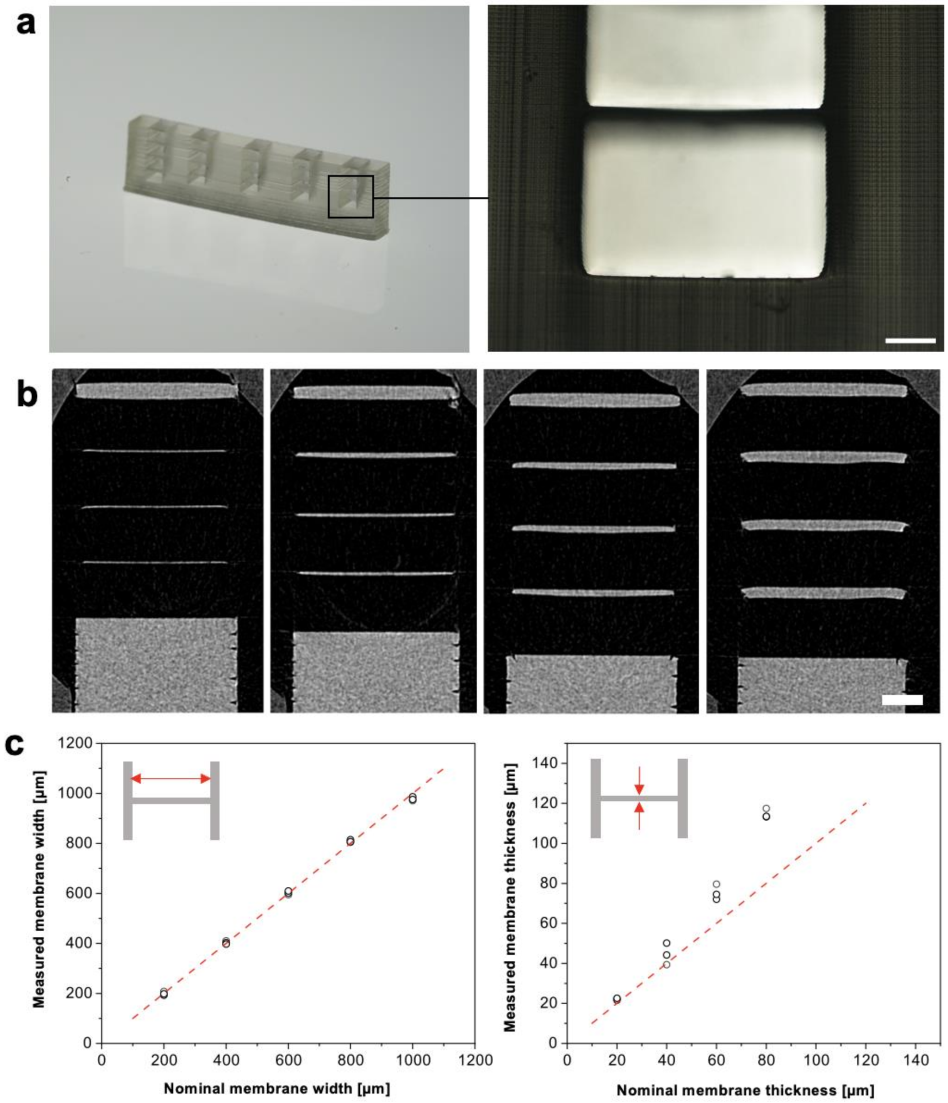


**Figure S8. *In situ* membrane 3D printing.** (a) Membrane calibrator device showing varying membrane thicknesses to assess the z-resolution, inset shows a 20 μm membrane. Scale bar = 200 μm. (b) μCT side views of 20, 40, 60, and 80 μm thick membranes. Scale bar = 500 μm. (c) Dimensional accuracy between design file and printed membrane widths and thickness across three unique devices. Dashed line shows the case where the nominal value = measured value.

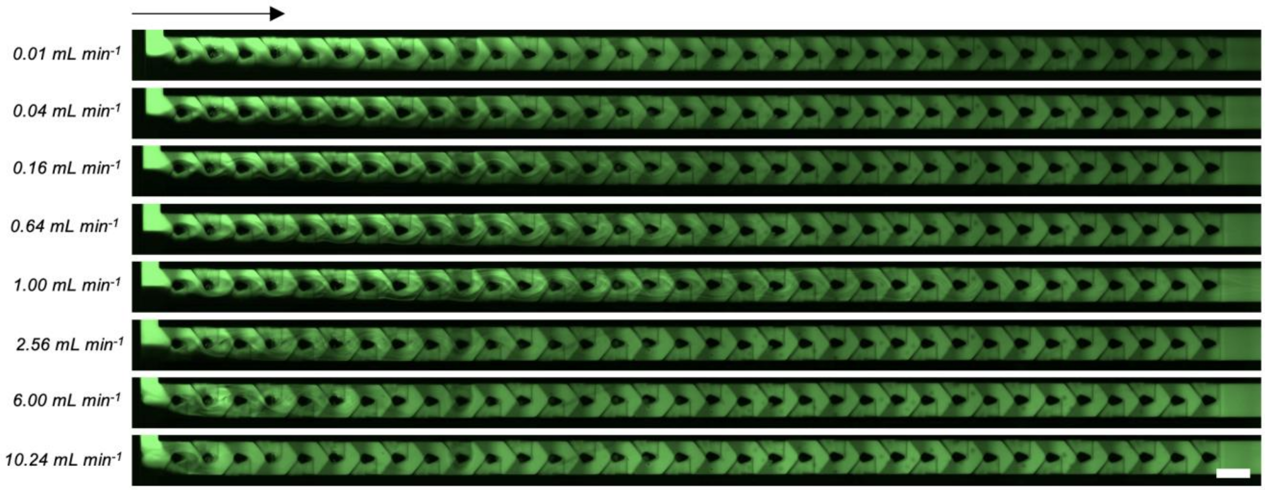


**Figure S9. Fluorescent images of microfluidic mixing at different volumetric flow rates.** Deionized water and 10 μM fluorescein solution were mixed over a range of flow rates from 0.01-10.24 mL min$^{-1}$ via fluorescence microscopy along the length of the channel. Arrow shows the direction of flow. Scale bar = 1 mm.

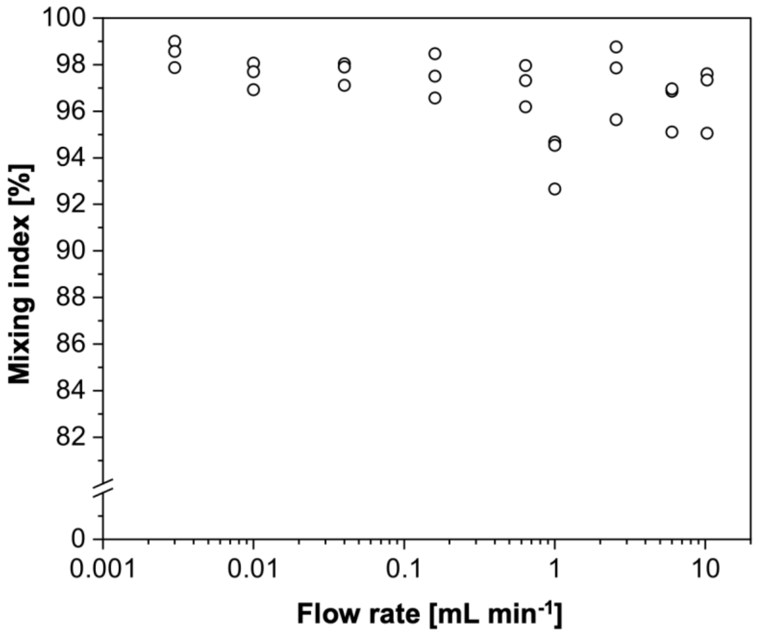


**Figure S10. Mixing indices**, shown for the mixing of 10 µM fluorescein and deionized water at different flow rates for three replicate micromixer devices.

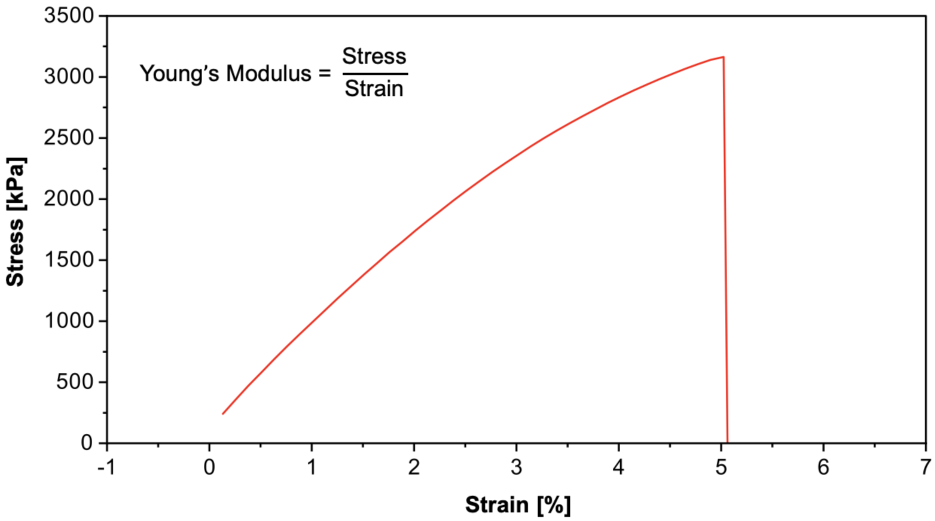


**Figure S11. Tensile testing of 3D printed parts using PLInk.** Application of a known load and observation of the measured extension gives a Young's modulus of 68 ± 3 MPa across three replicates.

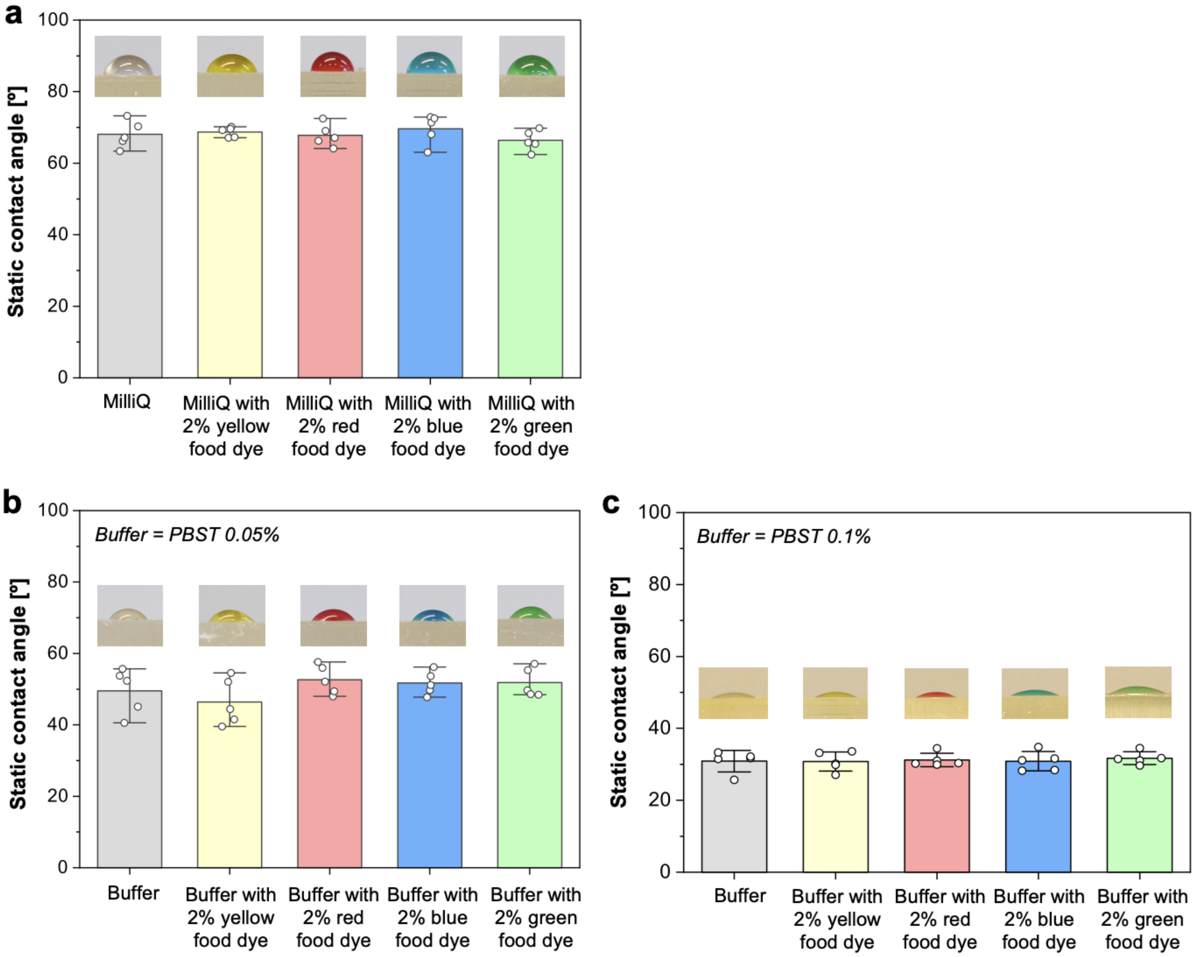


**Figure S12. Contact angle.** Contact angle of (a) water, (b) PBST 0.05%, and (c) PBST 0.1% ELISA buffer on 3D printed PLInk. Data shows mean ± STD across five replicates. Notably, this ink is inherently more hydrophilic than most commercial counterparts that we previously used for capillarics.[3, 4]

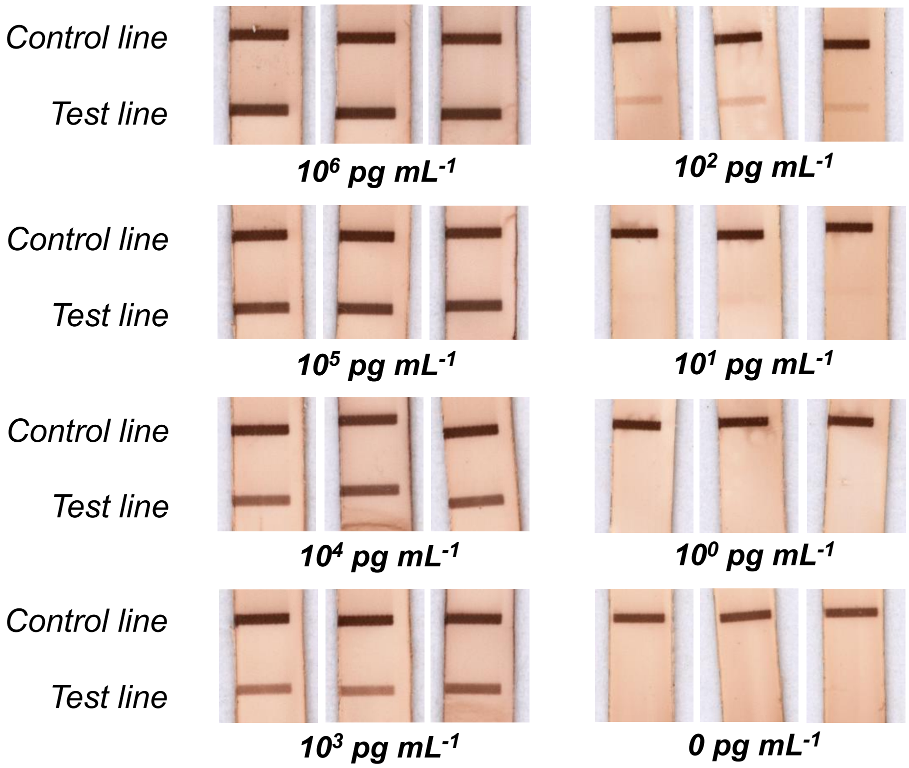


**Figure S13. Nitrocellulose membranes from LCD 3D printed ELISA-chip for IFN-γ detection.** Colorimetric readouts on a nitrocellulose membrane showing a control line and test line at various IFN-γ concentrations.

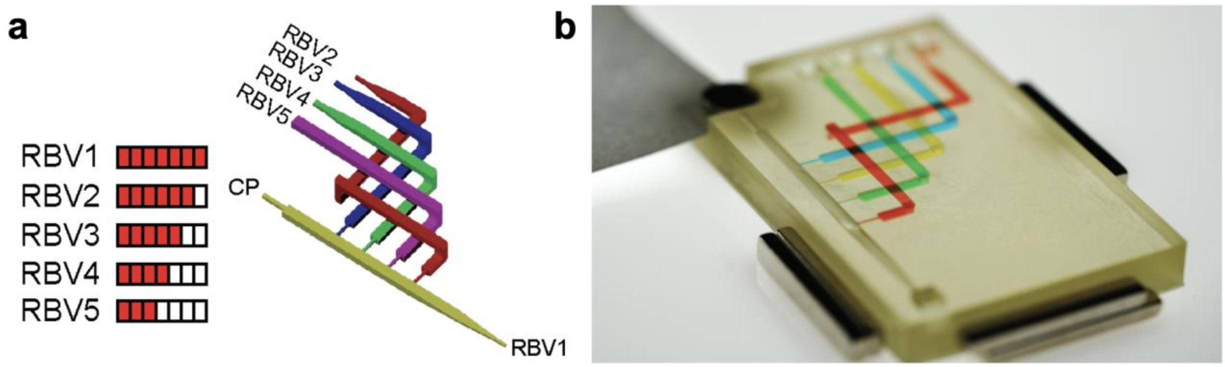


**Figure S14. Monolithic CC.** (a) Circuit design of an embedded CC device for the sequential delivery of four reagents based on their burst threshold pressures structurally encoded by capillary retention burst valves (RBVs) with decreasing cross-sections. The smallest cross-section RBV (i.e., the highest burst pressure threshold given by RBV1) was connected to the main channel, which was connected to a capillary pump (CP). (b) Image of 3D printed embedded CC with overlapping and weaving conduits.

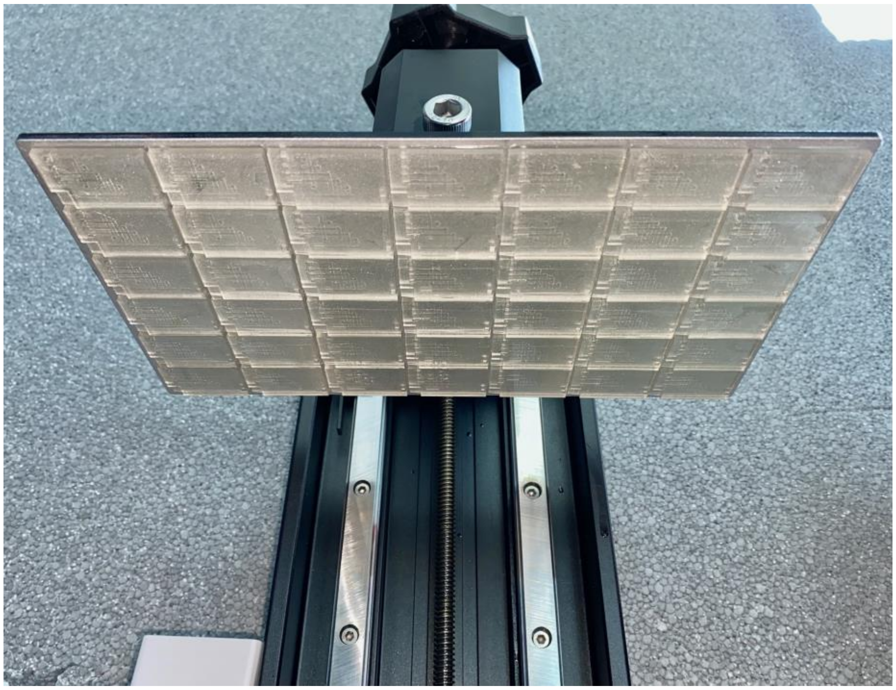

**Figure S15. Throughput manufacturing of monolithic CCs.** Using the Elegoo Saturn 2 8K LCD 3D printer with a nominal pixel size of $28.5 \times 28.5\ \mu m^2$, over ~33M pixels, and a build area of $225 \times 129\ mm^2$, 42 monolithic CCs were manufactured in <45 min.
